# Inhibitory actions of melanin-concentrating hormone in the lateral septum

**DOI:** 10.1101/2023.10.21.562777

**Authors:** Mikayla A Payant, C Duncan Spencer, Melissa J Chee

## Abstract

Melanin-concentrating hormone (MCH) neurons can coexpress several neuropeptides or neurotransmitters and send widespread projections throughout the brain. Notably, there is a dense cluster of nerve terminals from MCH neurons in the lateral septum (LS) that innervate LS cells by glutamate release. The LS is also a key region integrating stress- and anxiety-like behaviours that are also emerging roles of MCH neurons. However, it is not known if the MCH peptide acts within the LS or whether MCH target sites are localized. We analysed the projections from MCH neurons in male and female mice anteroposteriorly throughout the LS and found spatial overlap between the distribution pattern of MCH-immunoreactive (MCH-ir) fibers with MCH receptor *Mchr1* mRNA hybridization or MCHR1-ir cells. This overlap was most prominent along the ventral and lateral border of the rostral part of the LS (LSr). Most MCHR1-labeled LS neurons laid adjacent to passing MCH-ir fibers, but some MCH-ir varicosities directly contacted the soma or cilium of MCHR1-labeled LS neurons. We thus performed whole-cell patch-clamp recordings from MCHR1-rich LSr regions to determine if and how LS cells respond to MCH. Bath application of MCH to acute brain slices activated a bicuculline-sensitive chloride current that directly hyperpolarized LS cells. This MCH-mediated hyperpolarization was blocked by calphostin C and suggested that the inhibitory actions of MCH were mediated by protein kinase C-dependent activation of GABA_A_ receptors. Taken together, these findings defined potential hotspots within the LS that may elucidate the contributions of MCH to stress- or anxiety-related feeding behaviours.

**Key points:**

- ***RESEARCH QUESTION.*** Melanin-concentrating hormone (MCH) neurons have dense nerve terminals within the lateral septum (LS), a key region underlying stress- and anxiety-like behaviours that are emerging roles of the MCH system, but it is not known if the LS is a MCH target site.
- ***NEUROANATOMY.*** We found spatial overlap between MCH-immunoreactive fibers, *Mchr1* mRNA, and MCHR1 protein expression especially along the lateral border of the LS.
- ***ELECTROPHYSIOLOGY.*** Within MCHR1-rich regions, MCH directly inhibited LS cells by increasing a chloride conductance in a protein kinase C-dependent manner.
- ***SIGNIFICANCE.*** Electrophysiological MCH effects in brain slices have been elusive and even fewer have described the mechanisms of MCH action. Our findings demonstrated, to our knowledge, the first description of MCHR1 Gq-coupling in brain slices, which was previously predicted in cell or primary culture models only. Together, these findings defined hotspots and mechanistic underpinnings for MCH effects such as in stress- and anxiety-related behaviours.

## Introduction

Neurons that produce melanin-concentrating hormone (MCH) are found primarily within the lateral hypothalamic area (LHA) (Broberger *et al*., 1998; Broberger, 1999; Croizier *et al*., 2010; Beekly *et al*., 2020), but they can send widespread projections throughout the brain (Skofitsch *et al*., 1985; Bittencourt *et al*., 1992). MCH neurons can express additional neuropeptides (Harthoorn *et al*., 2005; Mickelsen *et al*., 2017) and neurotransmitters like GABA (Jego *et al*., 2013) and glutamate (Chee *et al*., 2015). MCH has well-established functions in energy balance (Qu *et al*., 1996; Shimada *et al*., 1998; Ludwig *et al*., 2001; Kokkotou *et al*., 2005; Pissios *et al*., 2006) and sleep (Verret *et al*., 2003; Ferreira *et al*., 2017), but recent findings have also elaborated on the roles of MCH for regulating stress (Kim & Han, 2016), motivation (Mul *et al*., 2011), and memory (Monzon et al., 1999; Adamantidis et al., 2005; Adamantidis and Lecea, 2009). The diverse functions of MCH thus implicate distinctive target sites for MCH.

MCH neurons strongly innervate the lateral septum (LS) via direct glutamatergic projections (Chee *et al*., 2015), but it is not known whether MCH plays a role in the LS. MCH immunoreactivity has been detected in the LS of the rat brain (Skofitsch *et al*., 1985; Bittencourt *et al*., 1992), but this has not been examined in detail for the mouse brain. In rats, immunohistochemical staining showed that the LS comprises moderate levels of MCH-immunoreactive (MCH-ir) fibers within the LS, with the highest level of immunoreactivity in the ventral part of the LS (Bittencourt *et al*., 1992). In addition to the presence of MCH-ir fibers, the expression of MCH receptors (MCHR) also aid in identifying the LS as a potential target site of MCH action. There are two known MCH receptors in the human brain, MCHR1 and MCHR2 (Hill *et al*., 2001), but only MCHR1 is present in the rodent brain (Tan *et al*., 2002). Similar to the widespread distribution of MCH-ir fibers, many brain regions can express *Mchr1* mRNA within the rat (Lembo *et al*., 1999; Saito *et al*., 2001) and mouse brain (Chee *et al*., 2013). Indeed, there is a moderate level of *Mchr1* mRNA in both the rat (Lembo *et al*., 1999; Saito *et al*., 2001) and mouse LS (Chee *et al*., 2013).

MCHR1 is a G-protein coupled receptor that can couple to G_i_- (Hawes *et al*., 2000), G_q_- (Hawes *et al*., 2000), or G_s_-protein-mediated pathways (Pissios *et al*., 2003). However, MCH action in the brain is largely inhibitory by hyperpolarizing the membrane and suppressing action potential firing (Gao, 2009), for example at the lateral hypothalamus (Rao *et al*., 2008), nucleus accumbens (Georgescu *et al*., 2005; Sears *et al*., 2010), or medial septal nucleus (Wu *et al*., 2009). In this study, we assessed the neuroanatomical and electrophysiological premise for MCH action in the LS and determined whether MCH could inhibit the activity of LS cells.

We described the distribution of MCH-ir fibers, *Mchr1* mRNA, and MCHR1 protein in the mouse LS, and we used these fiber and cell maps to guide patch-clamp recordings to identify putative sites and mechanisms of MCH action. As the MCH system (Mystkowski *et al*., 2000; Mogi *et al*., 2005; Rondini *et al*., 2007; Takase *et al*., 2014; Terrill *et al*., 2020; Teixeira *et al*., 2020) as well as the LS has been shown to be sexually dimorphic, we completed our analyses in the male and female brain but determined that there were no sex differences in the neuroanatomical and electrophysiological effects of MCH. We observed similar distribution patterns between MCH-ir, *Mchr1* mRNA, and MCHR1 protein throughout the entire rostrocaudal extent of the LS and found that MCH directly inhibited LS cells by recruiting protein kinase C (PKC) and activating a GABA_A_ receptor-mediated chloride conductance. These findings indicate that MCH can act in the LS to regulate neuron activity and suggest that the LS is an important projection site for MCH functions.

## Materials and Methods

The use of all animals has been approved by the Carleton University Animal Care Committee on Animal Use Protocol 110940 in accordance with guidelines provided by the Canadian Council on Animal Care. All C57BL/6J wild type mice (stock 000664; Jackson Laboratory, Bar Harbor, ME) were bred in house and maintained on a 12-hour light-dark cycle (22–24°C; 40–60% humidity). All mice were given *ad libitum* access to food (Teklad Global Diets 2014, Envigo, Mississauga, Canada) and water.

### Neuroanatomy

#### Tissue processing

Mice were anesthetized with an intraperitoneal injection (i.p.) of chloral hydrate (700 mg/kg; MilliporeSigma, Burlington, MA) prepared in sterile saline, transcardially perfused with cold (4°C) saline (0.9% NaCl), then followed by fixation with 10% formalin (VWR, Radnor, PA). The brain was extracted from the skull, post-fixed overnight in 10% formalin (24 hr, 4°C), and cryoprotected in phosphate buffered saline (PBS) containing 20% sucrose and 0.05% sodium azide (24 hr, 4°C). Mice whose brains were processed for MCHR1 immunohistochemistry were perfused with saline followed by 250 mL of 10% formalin. Brains were post-fixed in 20% sucrose dissolved in 10% formalin (4 hr, 4°C) then cryoprotected as above.

All brains were sliced into five series of 30 μm coronal sections using a freezing microtome (Spencer Lens Co., Buffalo, NY). Two tissue series remained free-floating in PBS-diluted formalin (comprising PBS-azide and formalin in a 9:1 ratio) prior to immunohistochemical staining for MCH or MCHR1. Three tissue series were mounted onto Fisherbrand Superfrost Plus Microscope Slides (Fischer Scientific, Waltham, MA) to use for *in situ* hybridization. One series, designated as probe tissue, was used for *Mchr1* hybridization. Two adjacent series served as a positive control and a negative control to the probe tissue. The negative control tissue was later used for Nissl staining to parcellate and define the neuroanatomical boundaries of each slice. After tissues were mounted, the glass slides were air dried at room temperature (RT, 20–23°C; 1 hr), and then at −20°C (30 min) before being stored at −80°C.

#### Single-label immunohistochemistry

To detect MCH immunoreactivity, the tissue was washed in six 5-min exchanges of PBS and pretreated with 10 mM sodium citrate for 5 min (75°C) followed by 0.3% hydrogen peroxide in PBS for 20 min (RT). Following three 10-min PBS exchanges, the tissue was then blocked with 3% normal donkey serum (Jackson ImmunoResearch Laboratories, Inc., West Grove, PA) dissolved in PBS with 0.25% Triton-X (PBT) and 0.05% sodium azide for 2 hr (NDS; RT). After blocking, the tissue was incubated with an anti-rabbit MCH antibody (1:2,000; kindly provided by Dr. E. Maratos-Flier, Beth Israel Deaconess Medical Center; RRID: AB_2314774; (Elias *et al*., 1998; Chee *et al*., 2013) overnight in NDS (RT). The following day, the tissue was washed six times in PBS (5 min each) then incubated with a biotinylated goat anti-rabbit antibody (1:500; Jackson ImmunoResearch Laboratories; RRID: AB_2337965) prepared in NDS for 1 hr (RT). The tissue was washed three times in PBS for 10 min each and treated with avidin biotin horseradish peroxidase (PK-6100, Vector Laboratories, Newark, CA) in PBT for 30 min (RT). Tissue was washed in three 10-min PBS exchanges and underwent tyramine signal amplification by treating with PBT comprising 0.005% hydrogen peroxide and 0.5% borate-buffered biotinylated (Sulfo-NHS-LC biotin; 21335, Thermo Fisher Scientific, Waltham, MA) tyramine (T90344, MilliporeSigma) for 20 min (RT). Following three 10-min washes in PBS, the tissue was incubated with an Alexa Fluor 647-conjugated streptavidin antibody (1:500; Jackson ImmunoResearch Laboratories; RRID: AB_ 2341101) and NeuroTrace 435/455 (1:50; N21479, Thermo Fisher Scientific) in NDS without sodium azide for 2 hr (RT). Slices were then mounted on SuperFrost Plus microscope slides and coverslipped with ProLong Diamond Antifade Mountant (Thermo Fisher Scientific).

#### Dual-label immunohistochemistry

To detect MCHR1 immunoreactivity, tissue was first washed in six 5-min PBS exchanges and pretreated with 0.3% hydrogen peroxide in PBS for 20 min (RT). Following a set of three 10-min washes, the tissue was blocked in NDS for 2 hr (RT) and then incubated in anti-rabbit MCHR1 antibody (1:3,000; Thermo Fisher Scientific; RRID: AB_2541682) prepared in NDS for 48 hr (4°C). The tissue was rinsed with six 5-min PBS exchanges and incubated with a biotinylated goat anti-rabbit antibody (1:5,000) in NDS without azide for 1 hr (RT). Following three 10-minute washes, the tissue was incubated in avidin biotin horseradish peroxidase in PBT for 30 min (RT). The tissue was washed three times (10 min each, RT) and underwent tyramine signal amplification. After washing the tissue three times with PBS (10 min each, RT), it was incubated with a Cy3-conjugated streptavidin antibody (1:200, RT; Jackson ImmunoResearch Laboratories; RRID: AB_2337244).

The tissue was then washed three times with PBS (10 min each) and incubated with an anti-rabbit NeuN antibody (1:2,000; MilliporeSigma; RRID: AB_2571567) in NDS overnight (RT). The following day, the tissue was washed in six 5-min PBS exchanges and incubated with a donkey anti-rabbit Alexa Fluor 488-conjugate (1:500; Thermo Fisher Scientific; RRID: AB_2535792) and NeuroTrace 435/455 (1:50) in NDS without azide. Finally, the tissue was washed for 2 hr in PBS (RT) prior to mounting on SuperFrost Plus slides and coverslipped with ProLong Diamond Antifade Mountant. This MCHR1 antibody has been previously validated for ciliary expression (Diniz et al., 2020) and we have determined that there is no MCHR1 staining in the LS of male or female MCHR1-knockout mice (data not shown).

#### Triple-label immunohistochemistry

To determine the proximity of MCHR1- and MCH-immunolabeling at NeuN-labeled cells, brain tissues were prepared using procedures for optimized MCHR1 labeling. Tissues were treated to label MCHR1 immunoreactivity, as described above, followed by tyramine signal amplification and treatment with an Alexa Fluor 647-conjugated streptavidin antibody (1:200; Jackson ImmunoResearch Laboratories; RRID: AB_ 2341101). After rinsing with three PBS exchanges (10 min each), they were immediately incubated anti-rabbit MCH (1:2,000; RRID: AB_2314774) and anti-mouse NeuN (1:1,000; HB6429, Hello Bio, Princeton, NJ) in NDS overnight (RT). After the tissues were washed in six 5-min PBS exchanges, they were incubated with an NDS cocktail comprising donkey anti-rabbit Alexa Fluor 568-conjugate (1:1,000; Thermo Fisher Scientific; RRID: AB_2534017) and donkey anti-mouse Alexa Fluor 488-conjugate (1:500; Thermo Fisher Scientific; RRID: AB_141607) for 2 hr at RT, rinsed with PBS, then mounted onto SuperFrost Plus slides and coverslipped with ProLong Diamond Antifade Mountant.

#### In situ hybridization

We optimized *in situ* hybridization procedures using a RNAscope Multiplex Fluorescent Reagent Kit v2 (Advanced Cell Diagnostics (ACD), Newark, CA) and manufacturer instructions for fixed-frozen mouse brain tissue (Document 323100-USM, ACD). To promote tissue adherence, slides were removed from storage at −80°C, baked at 37°C for 45 min, dehydrated in an ethanol gradient (50%, 70%, 100%; 5 min each), and then air-dried for 15 min (RT) immediately prior to the start of tissue treatments.

Tissue was rehydrated in PBS for 5 min (RT), pretreated with 5–8 drops of hydrogen peroxide (323110, ACD) for 10 min (RT), washed twice in distilled water for 1 min each, and submerged in 100% ethanol for 15 min (RT) to promote tissue adherence. The slides were then placed inside a steamer (Oster, Boca Raton, FL) using a coplin jar filled with preheated distilled water for 10 s (99°C) before transferring into the Target Retrieval Reagent (322000, ACD) for 5 min (99°C). Following two 15 s washes in distilled water (RT), the slides were dehydrated in 100% ethanol for 3 min, and then washed in three PBS exchanges (1 min each). A hydrophobic barrier was then drawn around each slide with an ImmEdge pen (Vector Laboratories), and the slides were dried overnight (RT). The following day, the slides were washed twice in PBS for 2 min and then placed in 10% formalin for 30 min (RT). Slides were then washed twice in PBS for 2 min and the tissue was treated with 5–8 drops of Protease Plus (322331, ACD) and incubated in a HybEZ oven (310010, ACD) at 40°C for 30 min. After protease treatment, the slides were washed with two exchanges of distilled water for 1 min each.

RNAscope probes for *Mm-Ppib* (313911, ACD), *Bacillus dapB* (320871, ACD), and *Mm-Mchr1* (317491, ACD) were designated for positive control, negative control, or experimental targeting, respectively, and were applied directly to the slides to cover the tissue. The tissue was hybridized for 2 hr at 40°C in the HybEZ oven, then washed with three fresh exchanges (2 min each; RT) of 1× Wash Buffer (310091, ACD). The hybridization signal was amplified by alternating incubations in AMP-1 (40°C, 30 min; 323110, ACD), AMP-2 (40°C, 30 min; 323110, ACD), and AMP-3 (40°C, 15 min; 323110, ACD) with two Wash Buffer washes (2 min each).

*Mchr1* hybridization was then labeled with Cyanine 3 (Cy3) by treating tissue with HRP-C1 (40°C, 15 min; 323110, ACD), washing the tissue twice in Wash Buffer for 2 min (RT), and incubating the tissue with TSA plus Cy3 (1:750; NEL44E001KT, PerkinElmer, Waltham, MA) in TSA Buffer (322809, ACD) for 30 min in the 40°C oven. Slides were then washed twice in Wash Buffer (2 min each, RT) and incubated with HRP Blocker (323110, ACD) in the oven at 40°C for 15 min.

Where applicable, the tissue underwent immunohistochemical staining to label MCH-immunoreactive fibers, as adapted from Mickelsen and colleagues (2019). The tissue was blocked with NDS without sodium azide and applied to each slide for 30 min (RT). After blocking, the tissue was incubated with an anti-rabbit MCH antibody (1:2,000; RRID: AB_2314774) for 1 hr (RT). The tissue was thoroughly rinsed with two exchanges in PBS (2 min each) then incubated with a donkey anti-rabbit Alexa Fluor 647 conjugate (1:500; ThermoFisher Scientific; RRID: AB_2536183) for 30 min (RT). After washing the slides twice in Wash Buffer for 2 min (RT), 4–6 drops of 4′,6-diamidino-2-phenylindole (DAPI; 323110, ACD) were applied for 30 s, and the slides were coverslipped using ProLong Diamond Antifade Mountant. Slides were dried in the dark overnight at RT and then stored at −20°C.

### Microscopy

All images were acquired using a Nikon Ti2-E inverted microscope (Nikon Instruments Inc., Mississauga, Canada) and processed using NIS-Elements Imaging Software (Nikon).

#### Confocal imaging

Tiled confocal images were acquired with a Nikon C2 confocal system using 405-nm, 488-nm, 561-nm, and 640-nm excitation lasers to visualize DAPI or NeuroTrace, Alexa Fluor 488, Cy3, and Alexa Fluor 647 fluorophores, respectively. Full brain overview images of DAPI-labeled nuclei from *Mchr1* stained slices were acquired using a 4× objective (0.20 numerical aperture). Higher magnification images of the LS used for analysis were imaged for DAPI/NeuroTrace, Alexa Fluor 488, Cy3, and/or Alexa Fluor 647 signals with a Plan Apochromat 10× objective (0.45 numerical aperture) or 20× objective (0.75 numerical aperture) at a single image plane and stitched with NIS-Elements Imaging Software. Where applicable, Z-stacks of 1 μm optical slices were acquired with a Plan Apochromat 40× objective (0.95 numerical aperture) or 60× objective (1.40 numerical aperture) and displayed as orthogonal *XY*, *XZ*, and *YZ* projections or projected by their maximum intensity values (NIS-Elements Imaging Software).

#### In situ hybridization signals

The negative control slices were imaged at a single image plane using the Plan Apochromat 10× objective and the 561-nm laser. The positive control *Ppib* hybridization signals were imaged to assess tissue and RNA quality. Images of all the sections containing the LS for both negative control and experimental probe series were acquired using the same settings to ensure that any differences observed between sections were not due to a difference in magnification, scan area, laser power, or gain. Tiled images of the LS from each probe section were then acquired at 10× magnification using the 405-nm, and 561-nm lasers to image DAPI- and *Mchr1*-labeling. Images were saved and exported such that the different channels could be toggled on and off to allow visualization of individual channels.

#### Brightfield imaging

Large field-of-view images of Nissl-stained tissue were viewed and imaged using a CF160 Plan Apochromat 10× objective lens and acquired with a DS-Ri2 colour camera (Nikon). Shading correction was applied during image acquisition to adjust for illumination inconsistences at the edge of each image tile. The tiled images were stitched with NIS-Elements Imaging Software.

### Image analysis

#### Plane-of-section analysis

To assess the neuroanatomical distribution of MCH-ir fibers and MCHR1 protein, we used unique cytoarchitectural features seen in tiled, confocal photomicrographs of NeuroTrace-staining to parcellate and draw boundaries corresponding to brain regions defined in the *Allen Reference Atlas* (*ARA*; Dong, 2008) (Supporting Figure 1A*i*, C*i*).

We used tiled, brightfield photomicrographs of Nissl-staining to parcellate tissue used to analyse *Mchr1* hybridization signals (Supporting Figure 1B*i*). Following confocal imaging, coverslipped tissue that served as the negative control was soaked in PBS overnight (RT) until the coverslip slid off. The exposed brain tissue was then treated for Nissl staining, as previously described (Negishi *et al*., 2020; Bono *et al*., 2022). Where necessary, DAPI-labeled overview images were aligned with parcellated images of the Nissl-stained tissue. Confocal images of *Mchr1* mRNA hybridization signal in the LS were imported into Adobe Illustrator 2021 (Adobe Inc., San Jose, CA) and aligned to DAPI-labeled overview images. White matter, ventricles, blood vessels, and other easily identifiable landmarks were used to ensure slices were properly aligned.

All parcellations were drawn in Illustrator using an Intuos graphic tablet (Wacom, Kazo, Japan) with reference to nomenclature and atlas levels provided by the *ARA*.

#### Fiber density

Confocal images of MCH immunoreactivity in the LS were visualized by Alexa Fluor 647 emission. MCH-ir axon fibers and varicosities were traced in a new layer within Illustrator using an Intuos graphic tablet (Supportingfigure 1A*ii*). Fiber tracing was restricted to the LS only and then mapped to *ARA* brain templates (see *Mapping* description below). For each atlas level, another layer was added to the Illustrator file so that a filled shape can be drawn to encompass each LS subregion and the entire LS area. A clipping mask of this filled shape was then applied to isolate, as separate image files, the filled shape of the total LS area, filled shape of each subregion, mapped fiber tracing in the full LS, and mapped fiber tracing in each subregion. The images were then analysed in MATLAB (MathWorks, Natick, MA) to determine the total number of pixels encompassing the subregions or entire LS area (*pixels_total_*) and the number of pixels occupied by the fiber tracings (*pixels_fibers_*). As the LS is a heterogenous three-dimensional structure, we analysed fiber density at all LS levels (Risold and Swanson, 1997a; Risold and Swanson, 1997b). The density of MCH-ir fibers at each *ARA* level (*D*) was expressed on a ratio scale as: *D = 100* (*pixels_fibers_/pixels_total_*) to capture nuanced changes in fiber density throughout the rostrocaudal axis of each LS subregion.

**Figure 1.**
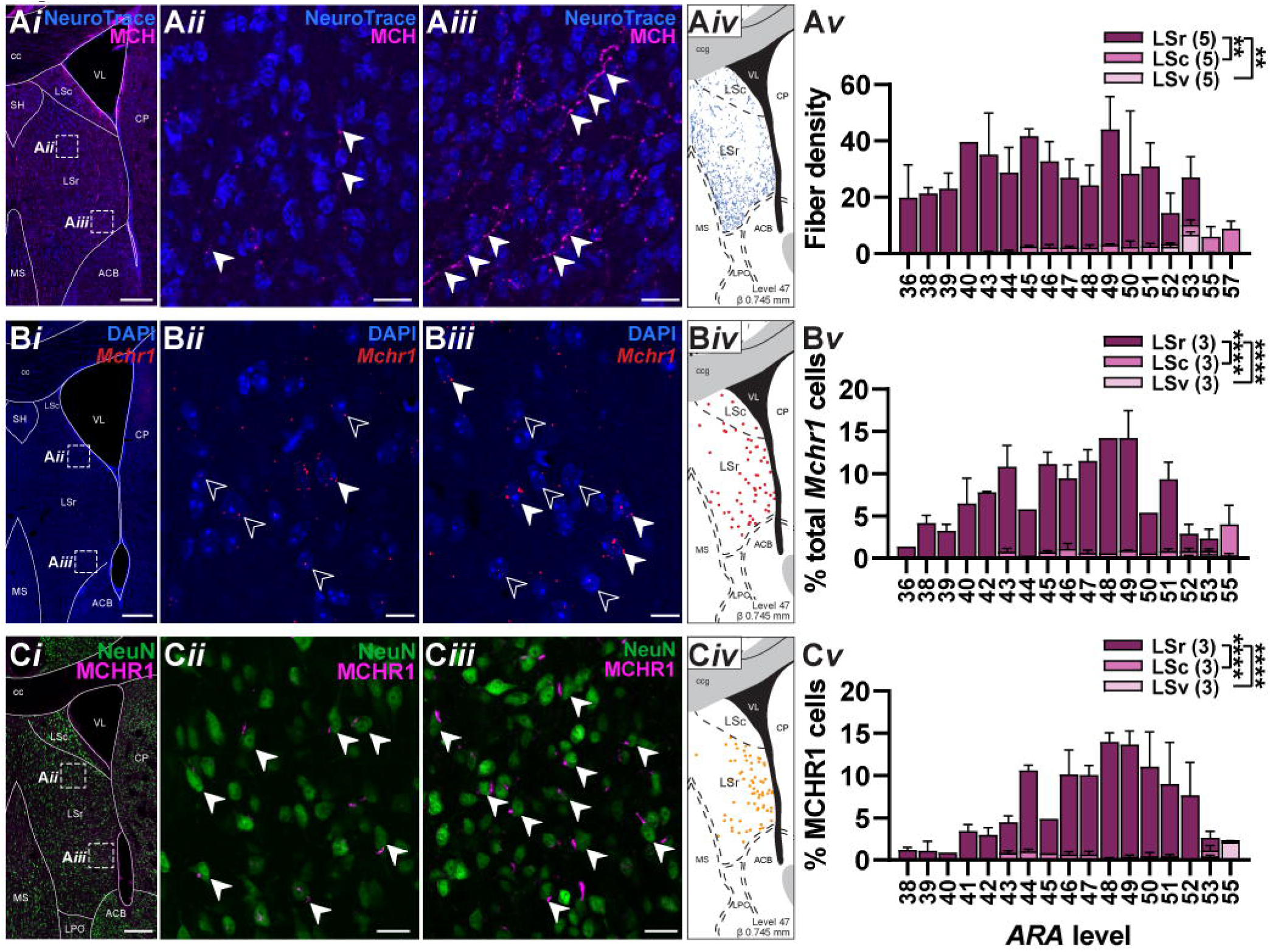
Relative expression of MCH-immunoreactive fibers, *Mchr1* mRNA, and MCHR1 protein throughout the LS. Representative confocal photomicrographs of MCH-immunoreactive (MCH-ir) fibers (arrowheads) amid NeuroTrace-labeled soma (***Ai***) in the dorsal (***Aii***) and ventral regions of the LS (***Aiii***). Coronal map of traced MCH-ir fibers in the LS (***Aiv***) at a representative *Allen Reference Atlas* level (*ARA*; Dong, 2008). Fiber density was expressed as the percent area covered by MCH-ir fibers at each *ARA* level in the rostral LS (LSr), caudal LS (LSc), and ventral LS (LSv; ***Av***). Representative confocal photomicrographs of *Mchr1* mRNA hybridization amid DAPI-labeled nuclei (***Bi***) showing 1–2 “dots” (open arrowhead; not included in subsequent analyses) or 3+ dots (white arrowhead) in the dorsal (***Bii***) and ventral regions of the LS (***Biii***). Only dots that surrounded a DAPI-labeled nucleus were included in our analyses. Coronal map of *Mchr1* hybridization distributed within the LS at a representative *ARA* level (***Biv***). Percent of *Mchr1* cells (comprising 3+ dots) at the LSr, LSc, and LSv of each *ARA* level was relative to the total number of *Mchr1* cells per brain (***Bv***). Representative confocal photomicrographs of MCHR1 immunoreactivity on the primary cilium (arrowhead) of NeuN-immunoreactive neurons (***Ci***) in the dorsal (***Cii***) and ventral regions of the LS (***Ciii***). Coronal map of MCHR1-expressing cells at a representative *ARA* level (***Civ***). Percent of total MCHR1 cells in the LSr, LSc, and LSv at each *ARA* level (***Cv***). Only *ARA* levels captured by our dataset were included. Scale bar: 200 μm (***Ai***, ***Bi***, ***Ci***), 25 μm (***Aii***, ***Aiii***), 10 μm (***Bii***, ***Biii***), 20 μm (**C*ii***, **C*iii***). Two-way mixed-effect model ANOVA with Tukey post-hoc testing: ** p < 0.01, **** p <0.0001. ACB, nucleus accumbens; cc, corpus callosum; CP, caudoputamen; LSc, lateral septal nucleus, caudal part; LSr, lateral septal nucleus, rostral part; MS, medial septal nucleus; SH, septohippocampal nucleus; VL, lateral ventricle.

#### Quantification of Mchr1 mRNA expression

Representative images from negative control tissue, corresponding to each probe slice, were adjusted using lookup table values (LUTs; NIS Elements) until the image appeared black to eliminate background fluorescence from any *dapB* hybridization. This set of LUTs were averaged and applied to images of *Mchr1* hybridization signals to subtract background fluorescence resulting from non-specific binding.

*Mchr1* hybridization was visualized by Cy3 fluorescence and appeared as punctate red dots, which were far fewer after background correction. Only dots colocalizing to a DAPI-stained nuclei were included in our analyses (Supporting Figure 1B*ii*). A DAPI-stained nucleus colocalizing with clusters of 3+ red dots were labeled as a *Mchr1*-expressing neuron and marked by a red-filled circle (Supporting Figure 1B*ii*). In the event that mRNA dots appeared between two DAPI-labeled nuclei, only one cell would be reported, thus it is possible that we are underestimating the number of *Mchr1* cells available in the LS. We counted the number of red-filled circles within the LS of each available brain slice.

#### Quantification of MCHR1 protein expression

Neuronal MCHR1 expression was counted from confocal images of NeuroTrace-labelled cells, ciliary MCHR1, and NeuN immunoreactivity visualized by NeuroTrace 435/455, Cy3, and Alexa Fluor 488 fluorescence, respectively. An orange-filled circle (Illustrator) was placed over NeuroTrace and NeuN-ir neurons marked by an MCHR1-ir primary cilium (Supporting Figure 1C*ii*). The number of circles were quantified within the LS of each available brain slice.

#### Mapping

The fiber tracings and filled circle labels were kept in individual layers of the Adobe Illustrator file so that each layer could be easily separated and mapped onto the corresponding level of the *ARA* template (Dong, 2008). The collection of fiber and circle labels was copied, resized, and adjusted so that the representation of the experimental LS fit the shape of the LS shown in the atlas reference template. In this way, neurons were mapped to their correct position relative to the unique neuroanatomical boundaries specific to the animal, despite physical differences unique to the animal (such as size and shape of brain regions). Individual subregions of the LS were mapped one-by-one to maintain accuracy in relative position and distribution of fibers and neurons (Supporting Figure 1A*iii*, B*iii*, C*iii*).

#### Appositions

Direct physical contact between fiber and soma or cilia was assessed using consecutive confocal Z-stack slices. Fiber contacts were referred to as appositions to the membrane where no visible space appeared between the fiber and cell membrane along the orthogonal *XZ* and *YZ* projections (Krimer *et al*., 1997; Lambe *et al*., 2000; Bouyer & Simerly, 2013). Contacts were determined at a physical zoom magnification of 2400× or greater, which permitted the detection of at least 0.4 μm gaps.

### Electrophysiology

#### Slice preparation

Mice were anesthetized with an injection of chloral hydrate (700 mg/kg, i.p.) and transcardially perfused with a carbogenated (95% O_2_, 5% CO_2_), ice cold artificial cerebrospinal fluid (ACSF) solution containing (in mM) 118 NaCl, 3 KCl, 1.3 MgSO_4_, 1.4 NaH_2_PO_4_, 5 MgCl_2_, 10 glucose, 26 NaHCO_3_, 0.5 CaCl_2_ (300 mOsm/L). The brain was removed from the skull and sliced at 250 μm using a vibrating microtome (VT1000s, Leica Biosystems, Buffalo Grove, IL) in cold, carbogenated ACSF. Slices containing the LS were transferred to glucose-based ASCF containing (in mM) 124 NaCl, 3 KCl, 1.3 MgSO_4_, 1.4 NaH_2_PO_4_, 10 glucose, 26 NaHCO_3_, 2.5 CaCl_2_ (300 mOsm/L) for 10 min (37°C) and then allowed to recover at RT for at least one hour prior to slice recording.

#### Slice recording

Slices containing the LS were bisected and transferred to the recording chamber where they were continuously perfused with carbogenated, glucose-based ACSF (31°C). Slice recordings were performed on three separate electrophysiology rigs. Cells were visualized with infrared differential interference contrast microscopy at 40× magnification on either an Examiner.A1 microscope (Zeiss, Oberkochen, Germany) equipped with an AxioCam camera (Zeiss) and Axiovision software (Zeiss), or with an Eclipse FN1 microscope (Nikon) equipped with a pco.panda 4.2 camera (Excelitas PCO GmbH, Kelheim, Germany) and NIS-Elements Imaging software (Nikon).

Whole-cell patch-clamp recordings were performed using borosilicate glass pipettes (7–9 MΩ) backfilled with a potassium-based internal pipette solution containing (in mM) 120 K-gluconate, 10 KCl, 10 HEPES, 1 MgCl_2_, 1 EGTA, 4 MgATP, 0.5 NaGTP, 10 phosphocreatine (290 mOsm/L, pH 7.24) to assess membrane properties, ionic conductances, and glutamatergic events. Internal pipette solution with an increased chloride concentration contained (in mM) 109 K-gluconate, 22 KCl, 10 HEPES, 1 MgCl_2_, 1 EGTA, 0.03 CaCl_2_, 4 MgATP, 0.5 NaGTP, 9 phosphocreatine (290 mOsm/L, pH 7.24). A cesium-based internal pipette solution used to record GABAergic events contained (in mM) 128 CsMS, 11 KCl, 10 HEPES, 0.1 CaCl_2_, 1 EGTA, 4 MgATP, 0.5 NaGTP (290 mOsm/L, pH 7.24). For recordings measuring membrane properties, 0.4% biocytin (Cayman Chemical, Ann Arbor, MI) was added to the internal pipette solution to allow for post-hoc immunohistochemical labeling and visualization of recorded cells. Recordings of electrical activity were generated using a MultiClamp 700B amplifier (Molecular Devices, San Jose, CA) and digitized by a Digidata 1440A (Molecular Devices) or using an Axopatch 200B amplifier (Molecular Devices) and digitized by a Digidata 1322A (Molecular Devices). All traces were acquired using pClamp 10.3 software (Molecular Devices) and filtered at 1 kHz.

#### Drug treatment

Following a baseline period of at least 5 min, MCH (3 μM; H-1482; Bachem, Torrance, CA) was bath applied into the recording chamber for approximately 5 min followed by a washout period in ACSF. Where applicable, tetrodotoxin (TTX; 500 nM; T-550, Alomone labs, Jerusalem, Israel), TC-MCH 7c (10 μM; 4365, Tocris, Toronto, Ontario, Canada), and bicuculline (30 μM; 14343, MilliporeSigma) were applied to the slice during the baseline period approximately 10 min prior to MCH application and maintained over the washout period. Calphostin C (100 nM; HB0160, Hello Bio Inc., Princeton, NJ) prepared and maintained in the dark until it was illuminated by a bright light within the slice recording chamber was applied to the slice for 20–30 min prior to MCH application. Antagonists were only added to cells that were hyperpolarized by a puff of MCH. All drugs were prepared from stock solution then dissolved into ACSF immediately prior to application.

#### Puff application

In experiments elucidating the membrane or intracellular mechanisms underlying the effects of MCH, we first delivered a short puff of MCH to a patched cell to identify those cells that responded with a reversible membrane hyperpolarization. To deliver the MCH puff, a second borosilicate glass “puff” pipette was filled with 3 μM MCH solution and lowered into the slice within 30–40 μm from the patched cell. A gradual positive pressure was manually applied to the puff pipette for 5–10 seconds until the MCH solution reached the patched cell.

#### Biocytin immunohistochemistry

Some brain slices used for electrophysiology recordings were post-fixed with 10% formalin to use for post-hoc immunohistochemical staining. The slices containing the biocytin-filled cells were rinsed in PBS (six 5-min washes), blocked in NDS (2 hours; RT), incubated with a streptavidin-conjugated Cy3 antibody (1:500) prepared in NDS (2 hours; RT), and then washed in PBS for 10 min. The slices were then washed with two more exchanges of PBS containing DAPI (1:2,000; Thermo Fisher Scientific) for 10 min. Brain slices were then mounted to Superfrost Plus microscope slides and coverslipped with ProLong Diamond Antifade Mountant.

### Experimental design and statistical analyses

#### Anatomical studies

Male and female mice wildtype mice (8–10 weeks) were used in a between-subject design to assess the distribution of MCH-ir fibers, *Mchr1* mRNA, and MCHR1 immunoreactivity. Comparisons between LS subregions or across LS levels were determined by two-way mixed model ANOVA with Tukey post-hoc testing, as not all LS levels can be captured in every brain sample.

#### Slice recording

Acute brain slices were prepared from male and female wildtype mice (38 male, 30 female) aged 5–23 weeks. Cells were recorded from two to three slices containing the LS and corresponding to Bregma 1.145–0.345 mm. Data sets included 1–4 cells per mouse.

#### Resting membrane potential (RMP)

Only neurons that exhibited a stable membrane potential (varied <5 mV) for 5 minutes prior to drug application were included in our data analyses. All voltages were corrected for a +15 mV liquid junction potential. For bath applications, RMP was sampled every 1 s using Clampfit 10.7 (Molecular Devices) and binned into 30 s increments. Control value was the mean RMP averaged over 1 min immediately prior to MCH application. The change in RMP (Δ RMP) was determined at the peak effect of MCH, which was within 4–8 min of MCH application and following washout 5–10 min later. In puff experiments, RMP was sampled every 500 ms and binned into 2-second increments, the Δ RMP elicited by MCH was sampled 10–25 s after the puff. Within-group designs comparing control, MCH, and washout conditions were analysed using a repeated measure one-way ANOVA with Tukey post-hoc testing. Comparisons of Δ RMP over time between two drug treatment conditions were analysed using a repeated measure two-way ANOVA. Comparisons of Δ RMP after or at peak effect of drug treatment were compared using a one-way ANOVA with Tukey post-hoc testing.

#### I–V curve

Ionic conductance was measured in voltage clamp from a holding potential (V_h_) of −75 mV. Descending 10 mV voltage steps (250 ms) were applied from −55 mV to −125 mV. The mean reversal potential (V_rev_) was averaged based on the V_rev_ for each cell, which was determined as the *x*-intercept calculated from a line equation where the slope is calculated from the −55 mV and −65 mV steps or from current values at two adjacent voltage steps where the current changes from a negative to a positive value, where applicable. The V_rev_ was compared to the theoretical equilibrium potential of the chloride ion (E_Cl_) using a one-sample *t* test. Between-group differences in net currents evoked following MCH application in the absence or presence of bicuculline were compared using repeated measures two-way ANOVA with Bonferroni post-hoc comparison.

#### Synaptic activity

Spontaneous (sIPSC) or miniature (mIPSC) inhibitory postsynaptic current events were recorded at V_h_ = –20 mV while excitatory post synaptic currents (sEPSC, mEPSC) were recorded at V_h_ = –75 mV. The IPSC and EPSC frequency were analysed using MiniAnalysis (Synaptosoft) and binned into 30-second increments. The control value was taken as the mean of a 1 min sample between 0 and 4.5 min prior to MCH application. The percent change in frequency and amplitude were determined at the peak effect of MCH between 2 and 9.5 min after the onset of MCH application. The washout was taken between 10 and 19 min after MCH application. Statistical significance was determined using a repeated measure one-way ANOVA with Tukey post-hoc testing.

We generated cumulative probability plots by pooling the amplitude and interevent intervals from 200 IPSC events or 50 EPSC events from each cell from baseline, MCH, and washout recording periods. Differences in the distribution of IPSC or EPSC amplitudes or interevent intervals in cumulative probability plots were analysed using the Kolmogorov-Smirnov *t* test.

#### Graphs and illustrations

All data graphs were generated using Prism 9 (GraphPad Software, San Diego, CA). Results were considered statistically significant at *p* < 0.05. Representative sample traces data were exported from Clampfit and plotted in Origin 2018 (OriginLab Corporation, Northampton, MA). Manuscript figures were assembled in Illustrator.

## Results

In order to identify potential sites of MCH action within the LS, we quantified the relative expression of MCH-ir fibers (Figure 1A), *Mchr1* mRNA (Figure 1B), and MCHR1 receptors (Figure 1C) in each subregion and level of the LS and then mapped their distribution throughout the rostrocaudal axis of the LS that spans 2.125 mm between *ARA* level (L) 36 and L57.

### Distribution of MCH-ir fibers throughout the LS

To maximize the detection of MCH-ir fibers, we performed our immunohistochemical stains using tyramide signal amplification. We then traced these fiber projections so that they can be mapped onto *ARA* templates with reference to Nissl-based parcellations and systematically examine the distribution of MCH-ir fibers throughout the LS. The density and pattern of MCH-ir fiber expression was comparable between males and females (Supporting Figure 2), so their datasets were combined to assess the overall MCH-ir fiber density across the LS.

The LS includes the rostral LS (LSr), caudal LS (LSc), and ventral LS (LSv). The LSr comprised the largest cytoarchitectural subdivision of the LS, and majority of MCH-ir fibers in the LS were found in the LSr (F(2, 12) = 10.33, p = 0.0025). Notably, MCH-ir fiber density was more abundant in the ventral than dorsal aspects of the LSr (Figure 1A*i*–*iii*). Near the peak *ARA* level of MCH-ir expression, there was a distinctive pattern of MCH-ir fibers that were more concentrated at the midline or along the medial LSr border adjacent to the medial septal nucleus, along the lateral LSr border adjacent to the lateral ventricle, and along the ventral LSr border abutting the nucleus accumbens, lateral preoptic area, or bed nuclei of the stria terminalis (Figure 1A*iv*; see Supporting Figure 2 for MCH-ir fiber maps at all *ARA* levels of the LS).

MCH-ir fiber density differed throughout the anteroposterior axis of the LS (F(16, 99) = 4.46, p < 0.0001), and MCH-ir fiber density in the LSr gradually increased by nearly two-fold at its peak between L45–L49 and then diminished posteriorly (Figure 1A*v*). The cytoarchitectural boundary of the LSc is dorsal to the LSr, begins around L44, and then persists throughout the LS. The LSc is a small LS subregion and comprised relatively few dispersed MCH-ir fibers within the overall LS (Figure 1A*v*). The LSv emerged posteriorly in the LS starting at L52, and the LSv contained a moderate and evenly distributed MCH-ir fiber density (Figure 1A*v*).

### Distribution of *Mchr1*-expressing LS cells

To determine if the LS expressed receptors for MCH, we used RNAscope to label *Mchr1* mRNA hybridization in the LS; positive staining for *Mchr1* mRNA appeared as punctate dots. We observed cells that contained 1–2 dots, and while even low amounts of mRNA may be translated to protein (Greer et al., 2016; Lipo et al., 2022), we only considered cells with 3+ dots to provide a conservative estimation of *Mchr1*-positive cells (Figure 1B). Similar to the distribution of MCH-ir fibers, *Mchr1* hybridization was more prominent in the LSr (F(2, 72) = 116.1, p < 0.0001), where *Mchr1* cells were most prevalent in the ventral than dorsal LSr (Figure 1B*i*–*iii*), where they tend to be along the lateral LSr borders (Figure 1B*iv*; see Supporting Figure 3 for representative maps of *Mchr1* hybridization at all *ARA* levels of the LS).

The distribution of *Mchr1*-expressing cells was similar between males and females (Supporting Figure 3). The number of *Mchr1* cells differed rostrocaudally within the LS (F(16, 72) = 3.47, p = 0.0001) and peaked at L49 (Figure 1B*v*). Of the identified *Mchr1*-expressing cells, 89% were found in the LSr and only 10% and 1% were located in the LSc and LSv, respectively. In the posterior levels at L52 and L53, there were few *Mchr1* cells in the LSr but there was an increasing proportion of *Mchr1* cells dorsally in the LSc.

### Distribution of MCHR1-expressing LS cells

To determine if *Mchr1* transcripts were translated to protein, we performed an immunohistochemical stain to label MCHR1 immunoreactivity. As MCHR1 is concentrated on the primary cilium of neurons (Diniz et al., 2020), we determined its colocalization to a NeuroTrace and/or NeuN-ir soma (Figure 1C). The vast majority (93%) of MCHR1 cells were in the LSr (F(2, 69) = 84.95, p < 0.001), and the pattern of MCHR1 immunoreactivity was similar to *Mchr1* mRNA hybridization. MCHR1-expressing cells clustered toward the lateral border and ventral half of the LSr, while the medial LSr bordering the medial septum had few MCHR1 cells at all levels of the LS (Figure 1C*i*–*iv*; see Supporting Figure 4 for representative maps of MCHR1-expressing cells at all *ARA* levels of the LS).

MCHR1-expressing cells were differentially distributed throughout the anteroposterior axis of the LS (F(16, 69) = 3.65, p < 0.0001) and peaked between L48 and L49, where there were 5-fold more MCHR1 cells than in the anterior or posterior LS (Figure 1C*v*). About 5% of MCHR1 cells were found in the LSc and were distributed across several *ARA* levels. The LSv comprised ∼2% of MCHR1 cells, which emerged posteriorly and were clustered at L55 (Figure 1C*v*). Interestingly, while 99.9% MCHR1-ir cilia in LSr and LSc colocalized to a NeuN-labeled soma, about 56% of MCHR1-ir cilia in the LSv did not colocalize to NeuN, though they were associated with a NeuroTrace-labeled cell.

### Proximity of MCH-ir fibers at MCHR1-expressing LS cells

MCH is known to reach its receptor and target site by volume transmission (Noble *et al*., 2018). However, as MCH fibers and MCHR1 are featured in similar LS regions dorsoventrally and rostrocaudally (Figure 1), we also assessed the proximity between MCH-ir fibers and *Mchr1* mRNA or MCHR1 protein expression. MCH-ir fibers were present around *Mchr1*-expressing cells in the ventral LSr (Figure 2A) and in some cases appeared to be in close contact with an *Mchr1*-expressing cell (Figure 2B). However, since *Mchr1* hybridization was localized to DAPI-labeled nuclei, it was not possible to assess if MCH-ir fibers formed direct appositions with a *Mchr1* cell. To determine if MCH fibers may come in direct contact with MCHR1-expressing LS cells, we performed a stain for MCHR1 protein, MCH-ir fibers, and the neuronal marker NeuN to mark the cell body and examined MCH fiber appositions on the soma or cilia of MCHR1-expressing cells. The majority of MCHR1-expressing LS cells (111 of 123) were not contacted by MCH-ir fibers either at their cell bodies or cilia (Figure 2C). Most MCHR1 cells were not immediately adjacent to visible MCH-ir fibers, and some MCH-ir varicosities came within 0.4 μm of MCHR1-labeled cilia or their affiliated cell body without making direct physical contact (Figure 2D). Interestingly, MCH-ir fibers directly contacted about 10% of MCHR1 LS cells examined, and these appositions may occur at the NeuN-labeled cell body (Figure 2E) or MCHR1-ir cilium (Figure 2F). These findings suggested that MCH is preferentially transmitted through diffusion from local MCH fibers in the LS, but MCH fibers may also be in direct contact with LS cells for localized actions.

**Figure 2.**
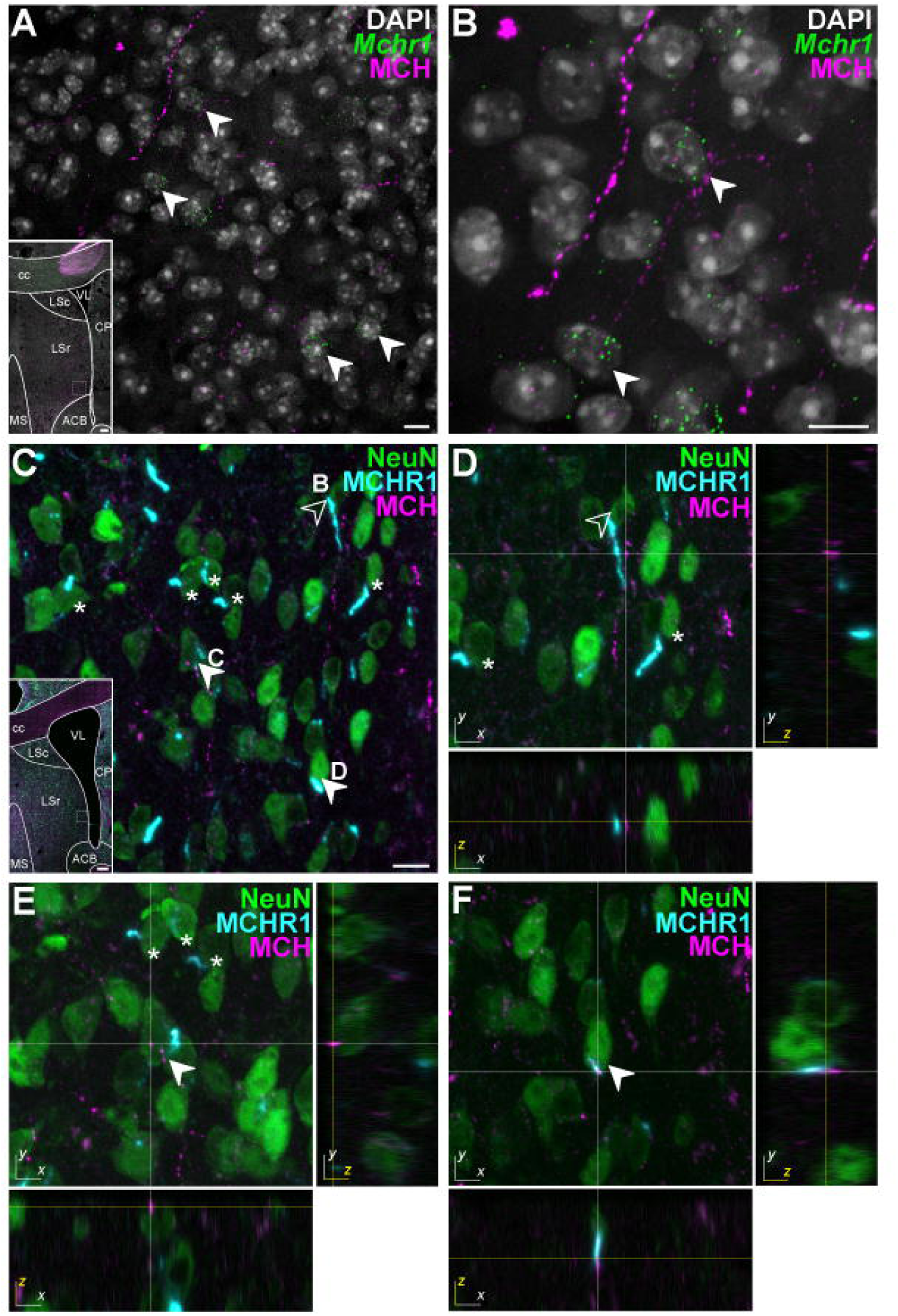
Proximity of MCH-immunoreactive fibers to MCHR1-expressing LS cells. Representative merged-channel confocal photomicrographs from the lateral and ventral LSr border (*inset*, dashed outlined area) of *Mchr1* mRNA on DAPI stained nuclei (white arrow) in relation to MCH-ir fibers (***A***) in close proximity to *Mchr1*-expressing cells (***B***). Representative merged-channel confocal photomicrographs from the lateral and ventral LSr border (*inset*, dashed outlined area) of MCHR1-ir cilia on NeuN-ir neurons in relation to MCH-ir fibers (***C***), which may be adjacent but relatively distant (asterisk) from MCHR1-ir LS cells. High magnification confocal photomicrographs with orthogonal projections in the *XY*-plane of MCH-ir varicosities in ***C*** that are closely associated but do not make physical contact (***D***, open arrowhead) or that form appositions (filled arrowhead) at the NeuN-ir soma (***E***) or associated MCHR1-ir cilia (***F***). Appositions were observed when no visible space was discerned between the MCH-ir varicosity and NeuN-ir soma or MCHR1-ir cilium in both the *XZ*- and *YZ*-plane at the same optical section (yellow line). Scale bars: 10 μm (***A***–***C***); 100 μm (*inset*, ***A***, ***C***); *X, Y*, *Z* axis 5 μm each (***B***–***D***). ACB, nucleus accumbens; cc, corpus callosum; CP, caudoputamen; LSc, lateral septal nucleus, caudal part; LSr, lateral septal nucleus, rostral part; LSv, lateral septal nucleus, ventral part; MS, medial septal nucleus; VL, lateral ventricle.

### MCH inhibited LS cells

As local diffusion may be the primary mode of MCH transmission within the LS, we hypothesized that substantive spatial overlap between the distribution of MCH-ir fibers and MCHR1 would define putative hotspots for MCH action. We found that the spatial overlap between the expression of MCH-ir fibers, *Mchr1* mRNA, and MCHR1 protein in the LS was most prominent toward the lateral and ventral borders of the LSr (Figure 3A), which formed our focus region for identifying MCH-responsive LS cells and defining the mechanisms of MCH action.

**Figure 3.**
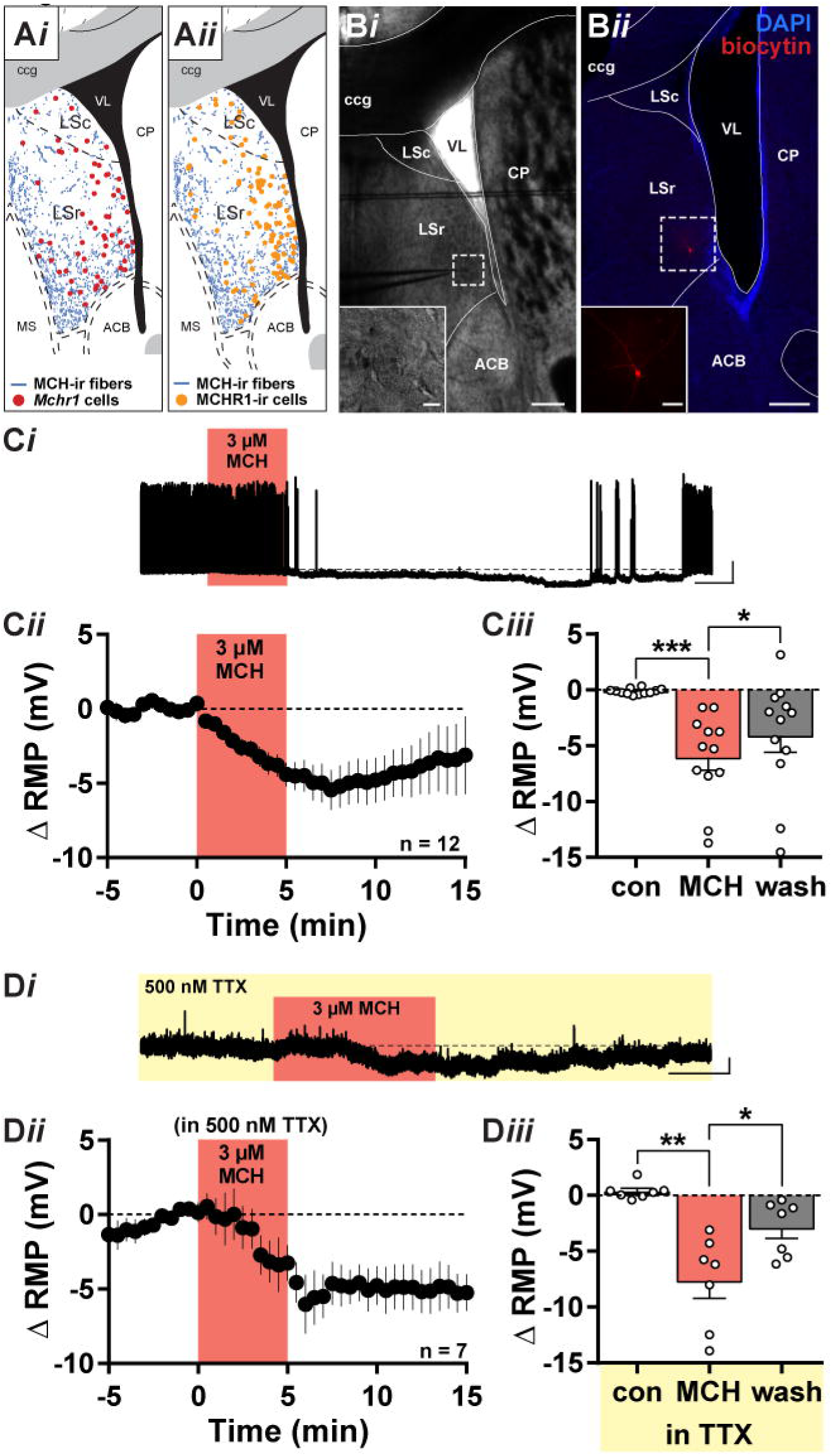
MCH directly hyperpolarized LS cells. Overlaid maps of MCH-immunoreactive (MCH-ir) fibers (blue) with low-*Mchr1* (pink circles; ***Ai***), high-*Mchr1* (red circles; ***Ai***), and MCHR1-ir cells (purple circles; ***Aii***) at *Allen Reference Atlas* Level 47 (Dong, 2008). Shape of the LS in acute brain slices guided whole-cell patch-clamp recordings (*inset*, ***Bi***) from cells near the ventrolateral border of the LSr (***Bi***). The position of the biocytin-filled recorded cell (*inset*, ***Bii***) was verified by post hoc staining (***Bii***). Representative sample trace of MCH-mediated hyperpolarization following bath application of 3 μM MCH (***Ci***). Time course of the MCH-mediated change in RMP (Δ RMP; ***Cii***) was summarized as the mean Δ RMP from each cell before MCH application (con), at the peak effect of MCH, and after MCH washout (wash) (***Ciii***). Representative sample trace of MCH-induced hyperpolarization in the presence of 500 nM TTX (***Di***). Time course of Δ RMP (***Dii***) was summarized as the mean Δ RMP from each cell before MCH application (con), at the peak effect of MCH, and after MCH washout (wash) (***Diii***). Scale bar: 200 µm (***Bi***, ***Bii***); 20 µm (***Bi*** *inset*); 50 μm (***Bii*** *inset*); 25 mV, 2 min (***Ci***); 4 mV, 2 min (***Di***). ACB, nucleus accumbens; ccg, corpus callosum, genu; CP, caudoputamen; LSc, lateral septal nucleus, caudal part; LSr, lateral septal nucleus, rostral part; MS, medial septal nucleus; VL, lateral ventricle.

We prepared acute brain slices containing the LS and performed whole-cell patch-clamp recordings from cells along the ventrolateral border of the LSr (Figure 3B). Bath application of MCH (3 µM) reversibly hyperpolarized the RMP of LS cells by −6.1 ± 1.1 mV (n = 12; F(2, 22) = 13.38, *p* = 0.002; Figure 3C). This MCH-mediated hyperpolarization was observed in about half of LSr neurons recorded from both male (7/12 cells) and female (5/9 cells) mice. We did not detect any differences in the magnitude of hyperpolarization between cells from male (−6.3 ± 1.5 mV, n = 7) or female mice (−5.6 ± 2.0, n = 5; t(10) = 0.29, *p* = 0.78), thus data from both male and female mice were combined.

To determine if MCH acts directly on LS cells, we pretreated the slice with TTX (500 nM) to block action potential-dependent activity. Subsequent co-application of MCH in the presence of TTX also hyperpolarized LS cells by −7.7 ± 1.5 mV (n = 7; F(2, 12) = 16.69, *p* = 0.002; Figure 3D), which indicated that MCH directly inhibited LS cells.

### MCH-mediated hyperpolarization is MCHR1-dependent

We screened for MCH-sensitive LS neurons by applying a short puff of MCH (Figure 4A*i*). We found that a single MCH puff (MCH_puff_) produced a small but reversible RMP hyperpolarization (−2.0 ± 0.4 mV, n = 8) that was sufficient to identify an MCH-sensitive neuron (Figure 4A*ii*). A second MCH_puff_ applied at least three minutes later also produced a similar (−1.6 ± 0.3 mV, n = 8) hyperpolarization (Figure 4A*ii*, A*iii*). By contrast, a puff application of bath ACSF did not alter the RMP (0.5 ± 0.3 mV, n = 9; F(2, 22) = 18.17, *p* < 0.0001; Figure 4A*ii*, A*iii*).

**Figure 4.**
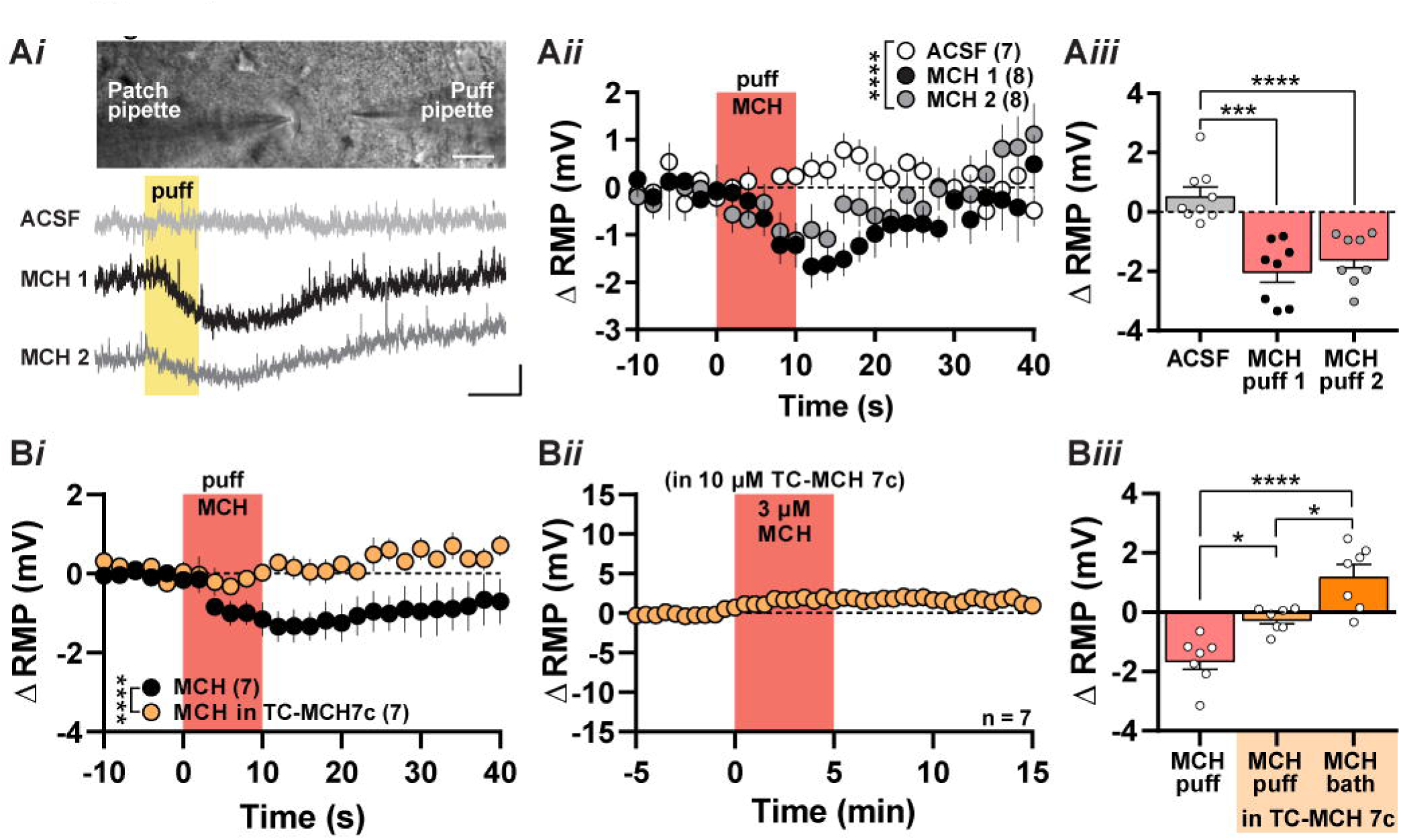
MCHR1-mediated hyperpolarization at LS cells. Brightfield photomicrograph showing the placement of a pipette for puff application during patch-clamp recording (*top*, ***Ai***). Representative sample trace of the change in resting membrane potential (Δ RMP) following a 10-second puff of ACSF or 3 μM MCH (MCH 1) followed by a subsequent MCH puff 3 minutes later (MCH 2) (*bottom*, ***Ai***). Comparison of Δ RMP following a puff of ACSF or 3 μM MCH over time (***Aii***) or immediately after the puff application (***Aiii***). Comparison of Δ RMP elicited in the presence or absence of the MCHR1 antagonist TC-MCH 7c (10 μM) by a 10-second puff of MCH (***Bi***), 5-minute bath application of MCH over time (***Bii***), or immediately after MCH application (***Biii***). Scale bar: 20 µm (*top*, ***Ai***) 2 mV, 10 s (*bottom*, ***Ai***).

To confirm that MCH-mediated hyperpolarization was occurring via the MCHR1 receptor, we applied MCH in the presence of the MCHR1 antagonist, TC-MCH 7c. After identifying an MCH-sensitive cell (MCH_puff_: −1.6 ± 0.3 mV, n = 7), we pretreated the slice with 10 µM TC-MCH 7c for 10–15 min. There was a main effect of the MCHR1 antagonist (F(2, 18) = 21.24; *p* < 0.0001), as a second MCH_puff_ applied in the presence of TC-MCH 7c no longer produced a hyperpolarization (−0.2 ± 0.2 mV, n = 7; *p* = 0.013; Figure 4B*i*, B*iii*). Similarly, MCH application via the slice chamber over a longer duration also did not elicit a membrane hyperpolarization in the presence of TC-MCH 7c (MCH_bath_: 1.2 ± 0.4 mV, n = 7; *p* < 0.0001; Figure 4B*ii*, B*iii*). These findings indicated that MCHR1 activation mediated the inhibitory effects of MCH.

### MCH activated a chloride channel to hyperpolarize LS cells

We next sought to determine the ionic basis of the MCH-mediated hyperpolarization. We compared the current–voltage (I–V) relationship of MCH-sensitive cells before and after a full bath application of MCH. We determined the I–V relationship of the cell by recording the steady-state current change elicited by each voltage step (Figure 5A*i*). MCH application elicited a membrane current with a V_rev_ of −63.7 ± 8.1 mV (n = 8), which corresponded to the predicted E_Cl_ (−63.1 mV) under our conditions (t(7) = 0.17, *p* = 0.87; Figure 5A*ii*). We then elevated the internal chloride concentration to determine if MCH-mediated inhibition was chloride-dependent. With an elevated chloride internal, MCH activated a smaller amplitude current (n=7; t(13) = 1.8, *p* = 0.047) with a V_rev_ of −48.1 ± 5.1 mV that corresponded with the new predicted E_Cl_ of −44.8 mV (t(6) = 0.64, *p* = 0.55; Figure 5A*ii*, A*iii*).

**Figure 5.**
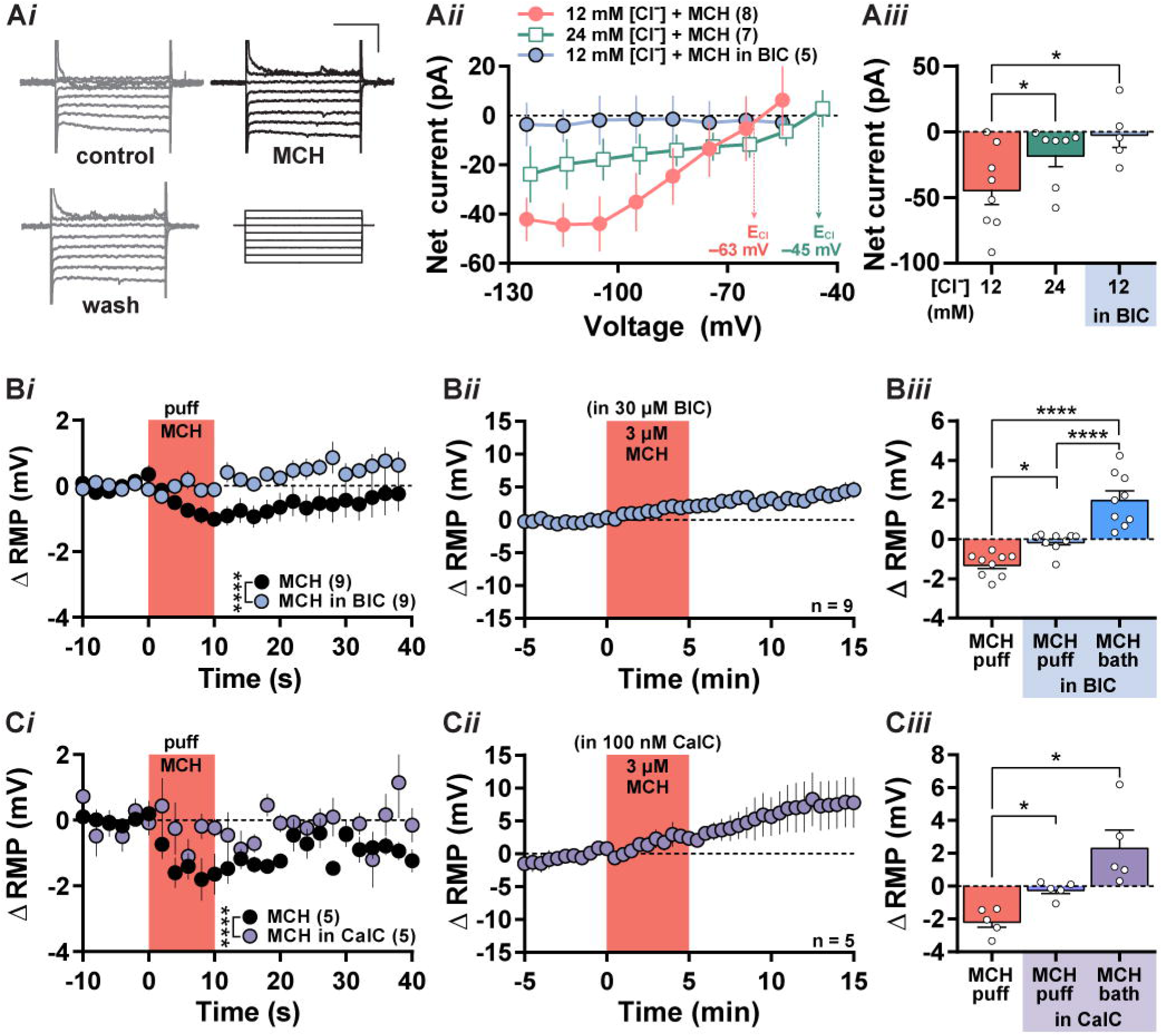
MCH-mediated activation of protein kinase C-dependent GABA_A_ receptors. Representative current output from MCH-sensitive cells immediately before (control), during a 3 μM MCH application, and following MCH washout (wash) in response to voltage steps (250 ms) applied at −10 mV increments from −55 mV to −125 mV (bottom right panel) (***Ai***). Comparison of the current–voltage relationship of net current elicited by 3 μM MCH in the absence (red filled circles) or presence of the GABA_A_ receptor antagonist bicuculline (BIC, 30 μM; blue filled circles) or with a high internal chloride concentration (green open squares) (***Aii***). Comparison of the change in resting membrane potential (Δ RMP) in the absence or presence of 30 μM BIC by a 10-second puff application of MCH over time (***Bi***), 5-minute bath application of MCH over time (***Bii***), or immediately after MCH application (***Biii***). Comparison of Δ RMP in the absence or presence of the protein kinase C inhibitor Calphostin C (CalC, 30 μM) by a 10-second puff application of MCH over time (***Ci***), 5-minute bath application of MCH over time (***Cii***), or immediately after MCH application (***Ciii***). Scale bar: 100 pA, 100 ms (***Ai***).

The net current elicited by MCH revealed an outward rectification at depolarizing potentials. These properties are consistent with that of ionotropic GABA_A_ receptor currents (Valeyev *et al*., 1999) expressed in the LS (Heldt & Ressler, 2007; Hörtnagl *et al*., 2013) and that contribute a tonic chloride conductance (Lee & Maguire, 2014). To determine if the MCH-mediated chloride current is related to the activation of GABA_A_ receptors, we next pretreated the slice with the GABA_A_ receptor antagonist bicuculline (30 μM), and subsequent MCH co-application did not evoke a change in membrane current (t(11) = 2.57, *p* = 0.026; Figure 5A*iii*). There was a significant interaction in the effect of bicuculline on the MCH-mediated current elicited across the voltage range (F(7, 77) = 7.38, *p* < 0.0001; Figure 5A*ii*) thus supporting a MCH-mediated activation of a GABA_A_ receptor.

To determine if this GABA_A_ receptor chloride conductance underlies the MCH-mediated hyperpolarization, we applied bicuculline to an MCH_puff_-sensitive cell (−1.3 ± 0.2 mV, n = 9). Bicuculline pretreatment abolished the inhibitory effects of MCH (F(2, 24) = 31.78, *p* < 0.0001), as neither a short MCH_puff_ (−0.1 ± 0.2 mV, n = 9; *p* = 0.027) nor prolonged MCH_bath_ application (2.0 ± 0.4 mV, n = 9; *p* < 0.0001) elicited a membrane hyperpolarization (Figure 5B). These findings indicated that a bicuculline-sensitive current underlies the inhibitory effect of MCH at LS cells and suggested that MCH may regulate the activation of a chloride conductance.

MCHR1 activation can couple to multiple intracellular G proteins, but most commonly to G_i_/G_o_ proteins or G_q_ proteins (Saito *et al*., 1999; Hawes *et al*., 2000). As PKC can be activated following MCHR1 stimulation (Pissios *et al*., 2003) and is linked to GABA_A_ receptor activation (Poisbeau *et al*., 1999) and increased chloride conductance via GABA_A_ receptors (Lin et al., 1996), we determined if MCH-mediated inhibition was linked to PKC activity. We first identified MCH-responsive cells (MCH_puff_: −2.1 ± 0.4 mV, n = 5) and pretreated the LS brain slice with 100 nM calphostin C, a PKC inhibitor. Interestingly, MCH application in the presence of calphostin C did not hyperpolarize the RMP (MCH_puff_: −0.2 ± 0.2, n = 5; *p* = 0.038) even when MCH was applied for an extended duration (MCH_bath_: 2.3 ± 1.1 mV, n = 5; *p* = 0.030). These findings indicated a main effect of calphostin C treatment (F(2, 8) = 11.92; *p* = 0.004; Figure 5C). Taken together, these results suggested that MCH activated a GABA_A_ receptor chloride current in a PKC-dependent manner.

### MCH did not inhibit GABAergic or glutamatergic input on LS cells

Previous MCH research showed that MCH can act via a presynaptic mechanism (Zheng *et al*., 2005), therefore we next determined if MCH also acts presynaptically in the LS. The LS comprises primarily GABAergic neurons with reciprocal connections to other LS cells (Gallagher & Hasuo, 1989; Sheehan *et al*., 2004; Zhao *et al*., 2013). We observed a high baseline frequency of sIPSC events (7.2 ± 3.1 Hz, n = 8) that was consistent with high GABAergic tone at LS cells (Carette et al., 2001; Figure 6A). MCH application produced a reversible rightward shift in the distribution of sIPSC interevent intervals (control vs. MCH: n = 8; D(8) = 0.27, *p* < 0.0001; MCH vs. wash: n = 8; D(8) = 0.30, *p* < 0.0001; Figure 6B, C) that reflected a 43% decrease in sIPSC frequency (baseline: 7.2 ± 3.1 Hz; MCH: 5.6 ± 2.7 Hz; wash: 6.4 ± 3.0 Hz; n = 8; F(2, 21) = 17.39, *p* < 0.0001; Figure 6C *inset*). In addition to decreasing sIPSC frequency, MCH produced a reversible leftward shift in the distribution of sIPSC event amplitudes (control vs. MCH: n = 8, D(8) = 0.23, *p* < 0.0001; MCH vs. wash: n = 8, D(8) = 0.30, *p* < 0.0001; Figure 6D) but did not decrease the mean amplitude of sIPSC events (control: 32.2 ± 5.0 pA; MCH: 29.8 ± 5.5 pA; wash: 31.9 ± 5.9 pA; n = 8; F(2, 21) = 0.67, *p* = 0.52; Figure 6D *inset*).

**Figure 6.**
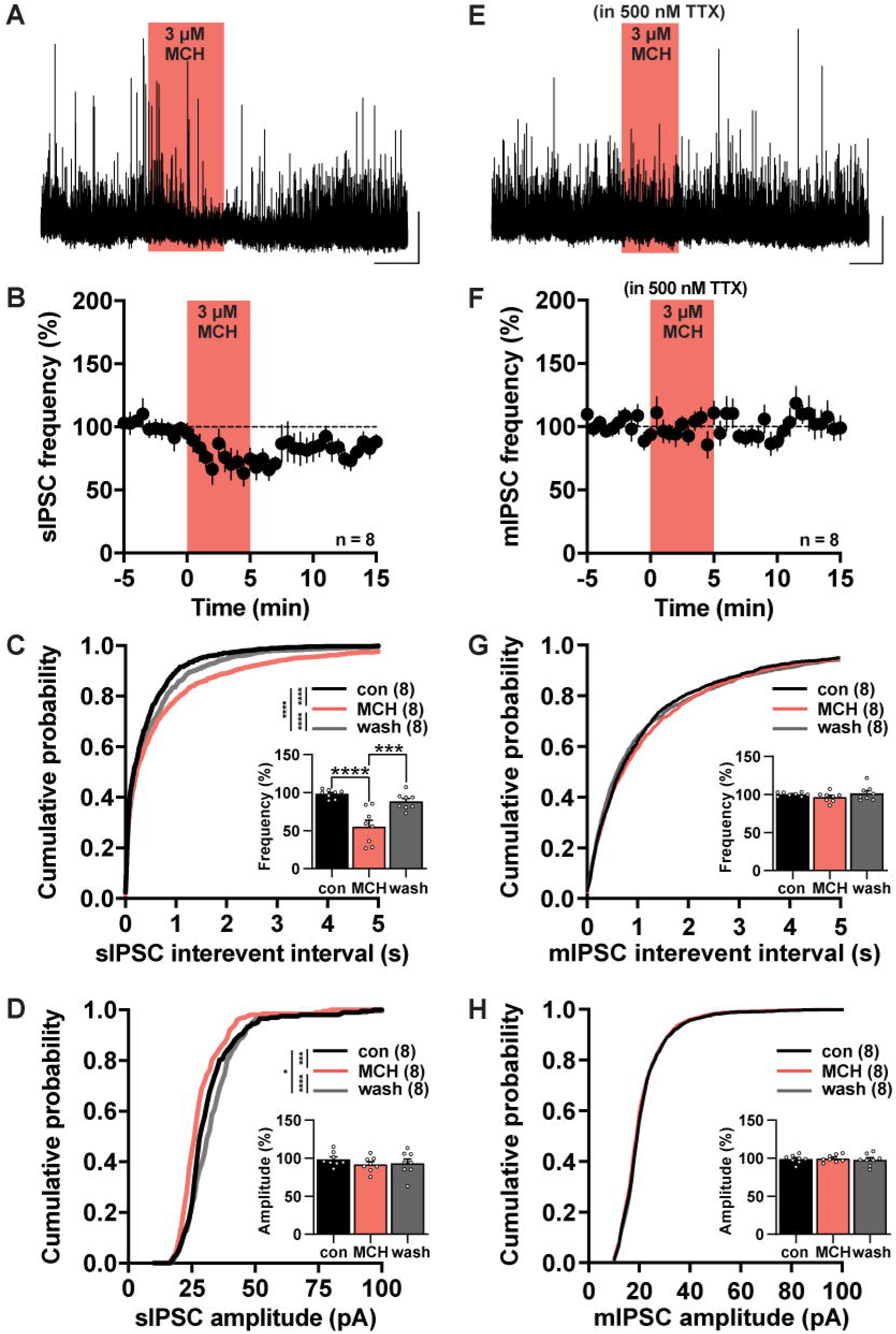
MCH suppressed spontaneous but not miniature IPSC events at LS cells. Representative sample traces of spontaneous IPSC (sIPSC; ***A***) and miniature IPSC (mIPSC) events recorded in the presence of 500 nM TTX (***E***) were summarized as the change in sIPSC (***B***) or mIPSC frequency over time (***F***). The cumulative distribution of sIPSC (***C***) and mIPSC interevent intervals (***G***) and amplitudes (***D***, ***H*)** were sampled immediately before (con), after MCH application, or following MCH washout (wash). Percent change in event frequency or amplitude was summarized in the *inset* (***C***, ***D***, ***G***, ***H***). Scale: 50 pA, 150 s (***A***, ***B***).

We pretreated the slice with 500 nM TTX to abolish activity-dependent synaptic transmission and recorded mIPSC events to determine if MCH would act on GABAergic presynaptic terminals (Figure 6E–H). Interestingly, co-application of MCH in TTX did not affect the distribution of interevent intervals (control vs MCH: n = 8; D(8) = 0.10; *p* = 0.27; Figure 6G) and, accordingly, did not change mIPSC frequency (control: 1.3 ± 0.4 Hz; MCH: 1.2 ± 0.3 Hz; wash: 1.3 ± 0.4 Hz; n = 8; F(2, 21) = 1.09, *p* = 0.35; Figure 6G *inset*). Likewise, MCH also did not change the distribution of mIPSC amplitudes (control vs MCH: n = 8, D(8) = 0.08, *p* = 0.47; Figure 6H) or mean mIPSC amplitudes (control: 21.2 ± 1.3 pA; MCH: 21.4 ± 1.4 pA; wash: 21.0 ± 1.3 pA; n = 8; F(2, 21) = 0.18, *p* = 0.84; Figure 6H *inset*). These findings suggested that MCH may not regulate presynaptic GABAergic inputs to LS cells.

The LS also receives glutamatergic input from different regions such as the hippocampus (Swanson & Cowan, 1977; Risold & Swanson, 1997a), which has also been shown to express MCHR1 (Saito *et al*., 1999). Therefore, we next examined if MCH can also change excitatory input to the LS. Spontaneous glutamatergic inputs to LS cells occur at a low frequency (1.3 ± 0.5 Hz, n = 9), and there was a slow rundown of sEPSC frequency over time (Figure 7A, B) that shifted the distribution of interevent intervals throughout the recording (control vs MCH: n = 9, D(9) = 0.54, *p* < 0.0001; control vs wash: n = 9, D(9) = 0.42, *p* = 0.0003; Figure 7C). Nonetheless, MCH did not alter sEPSC frequency (control: 1.3 ± 0.5 Hz; MCH: 1.3 ± 0.5 Hz; wash: 1.4 ± 0.6 Hz; n = 9; F(2, 16) = 0.60, *p* = 0.55; Figure 7C *inset*). MCH application also did not change the amplitude of sEPSC events (control: 15.5 ± 2.6 pA; MCH: 15.3 ± 2.3 pA; wash: 14.8 ± 2.0 pA; n = 9; F(2, 16) = 0.07, *p* = 0.91), which also reflected a gradual rundown over time (control vs MCH: n = 9, D(9) = 0.20, *p* = 0.21; control vs wash: n = 9, D(9) = 0.26, p = 0.05; Figure 7D).

**Figure 7.**
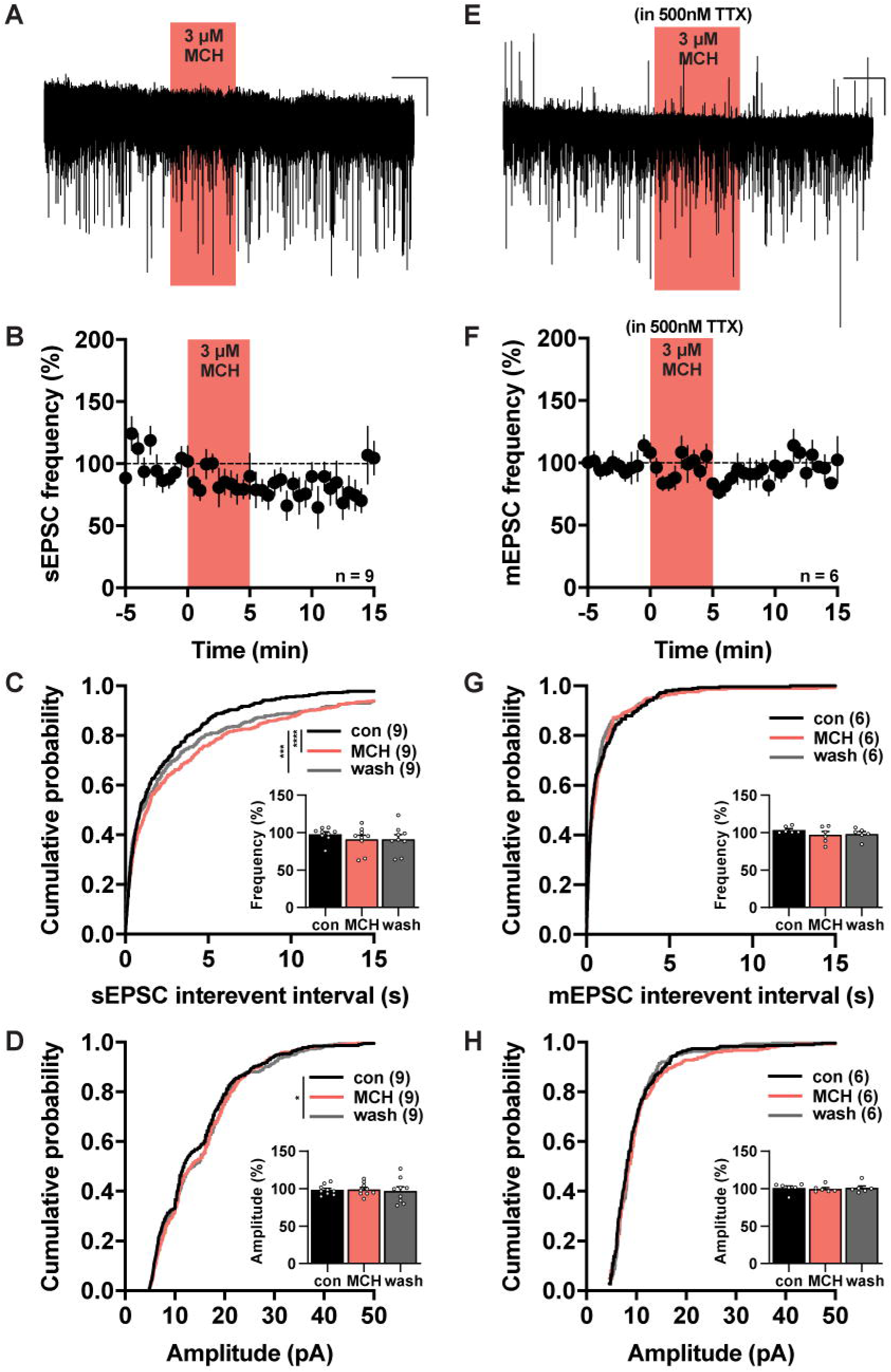
MCH did not regulate glutamatergic events at LS cells. Representative sample traces of spontaneous EPSC (sEPSC; ***A***) and miniature EPSC (mEPSC) events recorded in the presence of 500 nM TTX (***E***) were summarized as the change in sEPSC (***B***) or mEPSC frequency over time (***F***). The cumulative distribution of sEPSC (***C***) and mEPSC interevent intervals (***G***) and amplitudes (***D***, ***H***) were sampled immediately before (con), after MCH application, or following MCH washout (wash). Percent change in frequency or amplitude was summarized in the *inset* (***C***, ***D***, ***G***, ***H***). Scale: 10 pA, 150 s (***A***); 50 pA, 150 s (***B***).

In the presence of TTX, MCH had no effect on the frequency (control: 2.7 ± 1.0 Hz; MCH: 2.6 ± 1.0 Hz; wash: 2.6 ± 1.0 Hz; n = 6; F(2, 10) = 1.35; *p* = 0.32; Figure 7E–G) or the amplitude (control: 10.4 ± 1.1 pA; MCH: 10.1 ± 1.0 pA; wash: 10.2 ± 1.0 pA; n = 6; F(2, 10) = 0.06; *p* = 0.85; Figure 7H) of mEPSC events. Taken together, these findings suggested that MCH did not affect excitatory synaptic input.

## Discussion

MCHR1-expressing cells concentrated along the ventral and lateral boundaries of the LS and overlapped with the distribution pattern of MCH-ir fibers. We performed patch-clamp recordings from these MCHR1-rich hotspots and revealed a novel mechanism of MCH action in the LS. Pharmacological application of MCH directly hyperpolarized LS cells, but MCH did not alter synaptic input to LS cells. The inhibitory effects of MCH were MCHR1-dependent and mediated by an increased chloride current.

The coincident distribution pattern of *Mchr1* mRNA and MCHR1-ir cells in the LS implicated that these were MCH target sites. *Mchr1* hybridization signals can be seen throughout the LS but was most prominent in the LSr. This corresponded with *Mchr1* mRNA expression in the rat LS that was higher in the intermediate (Saito et al., 2001; i.e., mouse LSr) and ventral part of the LS (Bittencourt et al., 1992; i.e., mouse LSv) and lower in the dorsal part of the LS (Hervieu et al., 2000; i.e., mouse LSc). *Mchr1* hybridization signals reflected the presence of MCHR1 cells along the ventrolateral edge of the LSr anteriorly, as well as within the small LSv cluster posteriorly.

MCHR1-positive LS cells were distributed within or near MCH-ir fiber fields, which were also more abundant in the LSr than the LSc or LSv. Nerve terminals from MCH neurons are known to terminate in the LSr and form glutamatergic synapses to innervate LS cells (Chee *et al*., 2015). Interestingly, glutamatergic nerve endings from MCH neurons terminated in the dorsal zones of the LSr (Chee *et al*., 2015; Liu *et al*., 2022), where MCH-ir fibers were relatively sparse. Rather, MCH-ir fibers were prominent within the ventrolateral zones of the LSr where they spatially overlapped with MCHR1-expressing cells. MCH-ir fibers also extend into the medial zones of the LSr from the medial septum, but medially distributed MCH-ir fibers did not overlap with MCHR1-rich zones.

The proximity between MCH fibers and MCHR1-expressing LS cells implicated hotspots for MCHR1 activation. Most MCHR1 cells lie adjacent to but were not in contact with passing fibers and suggested that MCH is preferentially released extrasynaptically in the LS. MCH availability within the LS may also be a result of uptake from the lateral ventricle (Ruiz-Viroga *et al*., 2021) to then reach MCHR1 LS cells by volume transmission (Noble *et al*., 2018). This could be facilitated by MCHR1 expression along the primary cilium of neurons (Berbari *et al*., 2008; Diniz *et al*., 2020) to detect MCH in the extracellular space (Diniz *et al*., 2020). Furthermore, like in the nucleus accumbens (Sears *et al*., 2010) and nucleus of the solitary tract (Zheng *et al*., 2005) MCH-ir fibers can also be closely associated with LS cells, including MCHR1-expressing cells in the ventrolateral LS that were in direct physical contact with MCH-ir fibers. However, ultrastructural analyses would be required to determine if these contacts reflect active release sites. The functional contributions of local MCH release or MCH uptake from the ventricular space are not known (Ruiz-Viroga *et al*., 2021) but may influence behaviours that transpire over a longer timeframe. Meanwhile, local sources of MCH like those in direct contact with MCHR1 LS cells may mediate rapid, acute responses to environmental stressors.

Functional MCHR1 activation directly hyperpolarized and suppressed action potential firing of LS cells. While MCH can act presynaptically by inhibiting glutamatergic input (Zheng *et al*., 2005), it did not regulate either glutamatergic or GABAergic input to the LS. The inhibitory effect of MCH was mediated by a bicuculline-sensitive chloride current, which suggested the activation of ionotropic GABA_A_ receptors. While MCHR1 activation is known to couple to G_i_/G_o_-proteins (Gao, 2009), it can also couple to G_q_-proteins (Bächner *et al*., 1999; Pissios *et al*., 2003). MCHR1 coupling to the G_q_ pathway may activate chloride currents (Bächner *et al*., 1999) in a PKC-dependent manner (Pissios *et al*., 2003), as PKC activation can increase chloride influx via GABA_A_ receptors (Lin *et al*., 1994, 1996b). The postsynaptic mechanisms linked to MCH-mediated inhibition, including at the medial septal nucleus (Wu *et al*., 2009) and nucleus accumbens (Sears *et al*., 2010), have involved the activation of potassium channels, thus our findings implicate a novel chloride-mediated mechanism underlying the inhibitory actions of MCH.

The LS may integrate MCH roles that increase feeding (Qu *et al*., 1996; Rossi *et al*., 1997; Dilsiz *et al*., 2020) and anxiety-related behaviours (Smith et al., 2006; Dilsiz et al., 2020) by suppressing LS activity. Photostimulating GABAergic LS cells reduces feeding (Xu *et al*., 2019), and GABA_A_ and GABA_B_-mediated inhibition of LS cells can increase feeding (Gabriella et al., 2022; Calderwood et al., 2020). Since MCH can inhibit LS cells, MCH may thus increase feeding by downregulating GABAergic output from the LS. The LS is a notable region that regulates anxiety (Sheehan *et al*., 2004), and this may be ascribed to its afferent or efferent connections. MCH dampens LS activity, which can elicit anxiogenesis. For example, inhibiting hippocampal projections to the LS increased anxiety-like behaviours (Parfitt *et al*., 2017).

The LSc, LSr, and LSv designations by the *Allen Reference Atlas* are based on cytoarchitectonic parcellation (Dong, 2008), but LS divisions can also be informed by afferent and efferent connections that overlap with the distribution of MCHR1 cells (Risold and Swanson, 1997a; Risold and Swanson, 1997b). The ventrolateral LSr receives strong glutamatergic input from the ventral hippocampal CA1 (Risold and Swanson, 1997a) to suppress feeding and anxiety (Parfitt *et al*., 2017). Additionally, there is a concentration of cells expressing the type 2 corticotropin-releasing factor receptor (*Crfr2*) in the lateral LSr that innervate the anterior hypothalamic nucleus (Risold & Swanson, 1997a; Bang *et al*., 2022). Activating LS *Crfr2* terminals in the anterior hypothalamus promotes anxiety (Anthony *et al*., 2014).

The LS can also be divided into band-shaped domains informed by its chemoarchitecture (Risold and Swanson, 1997b). The distribution pattern of MCHR1-expressing LS cells corresponded to dorsolateral and ventrolateral LSr bands that overlap with enkephalin and urocortin immunoreactive fibers (Risold & Swanson, 1997b; Chen *et al*., 2011) anteriorly. These fibers have been implicated in feeding and anxiety-regulated behaviours at the LS. Enkephalin immunoreactivity is lower when food is less abundant and may reflect food availability (Kovacs *et al*., 2005). Enkephalin acts via the µ-opioid receptor in the LS (Mansour *et al*., 1994), and administration of a µ-opioid receptor agonist into the LSr increases feeding (Calderwood *et al*., 2020) and anxiety-like behaviour in mice (le Merrer *et al*., 2006). Urocortin activates CRFR2 receptors in the ventrolateral LS (Van Pett *et al*., 2000; Anthony *et al*., 2014) and can inhibit feeding during stress (Stengel & Taché, 2014). Urocortin administration into the LS inhibits feeding and increases anxiety-like behaviour (Wang & Kotz, 2002; Bakshi *et al*., 2007; Noguchi *et al*., 2013) and activation of *Crfr2* LS cells also promotes anxiety (Anthony *et al*., 2014). Interestingly, urocortin infusion into the LS can reduce orexin-mediated feeding (Wang & Kotz, 2002) and given that orexin and MCH have complimentary roles on feeding (Barson *et al*., 2013), interactions between urocortin and MCH may modulate feeding behaviour in response to stress.

MCH may also act via the LSv to mediate anxiety behaviours. MCHR1-expressing cells in the LSv may overlap with Substance P-immunoreactive fibers (Risold & Swanson, 1997b), and Substance P can have an anxiogenic effects in the LS (Gavioli *et al*., 1999, 2002; Ebner *et al*., 2008). The LSv receives glutamatergic input from the ventral hippocampus to promote coping mechanisms such as stress-induced grooming, which could help relieve feelings of stress and anxiety (Mu *et al*., 2020). Given the inhibitory actions of MCH on the LS, it may thus act via the LSv to dampen such neuronal coping mechanisms.

In conclusion, our findings showed that MCH inhibits the LS and suggests that MCH might converge with enkephalin and urocortin at the LSr, or with Substance P in the LSv, to fine-tune feeding and anxiety-related behaviours. We anticipated that the convergence of MCH fibers and MCHR1-expressing cells would assist in identifying hotspots underlying the action of MCH in the LS, and our electrophysiological recordings in these regions revealed a novel mechanism of MCH-mediated action that inhibited LS cells by increasing chloride conductance.

## Additional Information

### Data Availability

The data from this study are available upon reasonable request.

### Competing Interests

The authors have no conflicts of interest to declare.

### Author Contributions

Study conception and design: M.J.C. Acquisition, analysis, and interpretation of neuroanatomical datasets: M.A.P, C.D.S, M.J.C. Acquisition, analysis, and interpretation of electrophysiological datasets: M.A.P, M.J.C. Initial manuscript draft: M.A.P, C.D.S. Manuscript editing: M.A.P, M.J.C. All authors approved the final manuscript version, agree to be accountable for all aspects of the work, and agree that all authors that qualify for authorship are listed.

### Funding

This study was supported by Natural Sciences and Engineering Research Council of Canada (NSERC) Discovery Grant RGPIN-2017-06272 and Canadian Institutes of Health Research Project Grant 452284. M.A.P is supported by the NSERC Postgraduate Doctoral Scholarship and the Ontario Graduate Scholarship. C.D.S is supported by the NSERC Canadian Graduate Master’s Scholarship.

## Acknowledgements

The authors thank Dr. Ryan Chee for technical assistance writing MATLAB scripts.

## Supporting Information

Schematic of neuroanatomical analyses workflow (Supporting Figure 1). Detailed maps using *Allen Reference Atlas* brain templates showing the spatial distribution of MCH-ir fibers (Supporting Figure 2), *Mchr1* mRNA (Supporting Figure 3), and MCHR1 protein (Supporting Figure 4) expression rostrocaudally within the LS.

## Notes

### Competing Interest Statement

The authors have declared no competing interest.

## References

Adamantidis A, Thomas E, Foidart A, Tyhon A, Coumans B, Minet A, Tirelli E, Seutin V, Grisar T & Lakaye B (2005). Disrupting the melanin-concentrating hormone receptor 1 in mice leads to cognitive deficits and alterations of NMDA receptor function. Eur J Neurosci 21, 2837–2844.

Adamantidis A & de Lecea L (2009). A role for Melanin-Concentrating Hormone in learning and memory. Peptides (NY) 30, 2066–2070.

Anthony TE, Dee N, Bernard A, Lerchner W, Heintz N & Anderson DJ (2014). Control of stress-induced persistent anxiety by an extra-amygdala septohypothalamic circuit. Cell 156, 522–536.

Bächner D, Kreienkamp HJ, Weise C, Buck F & Richter D (1999). Identification of melanin concentrating hormone (MCH) as the natural ligand for the orphan somatostatin-like receptor 1 (SLC-1). FEBS Lett 457, 522–524.

Bakshi VP, Newman SM, Smith-roe S, Jochman KA & Kalin NH (2007). Stimulation of Lateral Septum CRF 2 Receptors Promotes Anorexia and Stress-Like BehaviorsL: Functional Homology to CRF 1 Receptors in Basolateral Amygdala. J Neurosci 27, 10568–10577.

Bang JY, Zhao J, Rahman M, St-Cyr S, McGowan PO & Kim JC (2022). Hippocampus-Anterior Hypothalamic Circuit Modulates Stress-Induced Endocrine and Behavioral Response. Front Neural Circuits; DOI: 10.3389/fncir.2022.894722.

Barson JR, Morganstern I & Leibowitz SF (2013). Complementary roles of orexin and melanin-concentrating hormone in feeding behavior. Int J Endocrinol 2013, 1–10.

Beekly BG, Frankel WC, Berg T, Allen SJ, Garcia-Galiano D, Vanini G & Elias CF (2020). Dissociated Pmch and Cre expression in lactating Pmch-Cre BAC transgenic mice. Front Neuroanat 14, 1–15.

Berbari NF, Johnson AD, Lewis JS, Askwith CC & Mykytyn K (2008). Identification of Ciliary Localization Sequences within the Third Intracellular Loop of G Protein-coupled Receptors. Mol Biol Cell 19, 1540–1547.

Bittencourt JC, Presse F, Arias C, Peto C, Vaughan J, Nahon J LL, Vale W & Sawchenko PE (1992). The melaninLconcentrating hormone system of the rat brain: An immunoL and hybridization histochemical characterization. J Comp Neurol 319, 218–245.

Bono BS, Ly NKK, Miller PA, Williams-Ikhenoba J, Dumiaty Y & Chee MJ (2022). Spatial distribution of beta-klotho mRNA in the mouse hypothalamus, hippocampal region, subiculum, and amygdala. J Comp Neurol 530, 1634–1657.

Bouyer K & Simerly RB (2013). Neonatal leptin exposure specifies innervation of presympathetic hypothalamic neurons and improves the metabolic status of leptin-deficient mice. J Neurosci 33, 840–851.

Broberger C (1999). Hypothalamic cocaine- and amphetamine-regulated transcript (CART) neurons: histochemical relationship to thyrotropin-releasing hormone, melanin-concentrating hormone, orexin/hypocretin and neuropeptide Y. Brain Res 848, 101–113.

Broberger C, de Lecea L, Sutcliffe JG, Hökfelt T & Hökfelt H (1998). Hypocretin/Orexin-and Melanin-Concentrating Hormone-Expressing Cells Form Distinct Populations in the Rodent Lateral Hypothalamus: Relationship to the Neuropeptide Y and Agouti Gene-Related Protein Systems. J Comp Neurol 402, 460–474.

Calderwood MT, Tseng A & Glenn Stanley B (2020). Lateral septum mu opioid receptors in stimulation of feeding. Brain Res 1734, 146648.

Carette B, Poulain P & Beauvillain JC (2001). Noradrenaline modulates GABA-mediated synaptic transmission in neurones of the mediolateral part of the guinea pig lateral septum via local circuits. Neurosci Res 39, 71–77.

Chee MJS, Arrigoni E & Maratos-Flier E (2015). Melanin-concentrating hormone neurons release glutamate for feedforward inhibition of the lateral septum. J Neurosci 35, 3644– 3651.

Chee MJS, Pissios P & Maratos-Flier E (2013). Neurochemical characterization of neurons expressing melanin-concentrating hormone receptor 1 in the mouse hypothalamus. J Comp Neurol 521, 2208–2234.

Chen P, Lin D, Giesler J & Li C (2011). Identification of urocortin 3 afferent projection to the ventromedial nucleus of the hypothalamus in rat brain. J Comp Neurol 519, 2023–2042.

Croizier S, Franchi-Bernard G, Colard C, Poncet F, la Roche A & Risold PY (2010). A comparative analysis shows morphofunctional differences between the rat and mouse melanin-concentrating hormone systems. PLoS One 5, e15471.

Dilsiz P, Aklan I, Sayar Atasoy N, Yavuz Y, Filiz G, Koksalar F, Ates T, Oncul M, Coban I, Ates Oz E, Cebecioglu U, Alp M, Yilmaz B & Atasoy D (2020). MCH Neuron Activity Is Sufficient for Reward and Reinforces Feeding. Neuroendocrinology 110, 258–270.

Diniz GB, Battagello DS, Klein MO, Bono BSM, Ferreira JGP, Motta-Teixeira LC, Duarte JCG, Presse F, Nahon JL, Adamantidis A, Chee MJ, Sita L v. & Bittencourt JC (2020). Ciliary melanin-concentrating hormone receptor 1 (MCHR1) is widely distributed in the murine CNS in a sex-independent manner. J Neurosci Res 0, 1–27.

Dong HW (2008). The Allen reference atlas: A digital color brain atlas of the C57BL/6J male mouse. John Wiley & Sons, Hoboken, NJ.

Ebner K, Muigg P, Singewald G & Singewald N (2008). Substance P in Stress and Anxiety NK-1 Receptor Antagonism Interacts with Key Brain Areas of the Stress Circuitry. Ann N Y Acad Sci 1144, 61–73.

Elias CF, Saper CB, Maratos-Flier E, Tritos NA, Lee C, Kelly J, Tatro JB, Huffman GE, Ollmann MM, Barsh GS, Sakurai T, Yanagisawa M & Elmquist JK (1998). Chemically defined projections linking the mediobasal hypothalamus and the lateral hypothalamic area. J Comp Neurol 402, 442–459.

Ferreira J, Bittencourt J & Adamantidis A (2017). Melanin-concentrating hormone and sleep. Curr Opin Neurobiol 44, 152–158.

Gabriella I, Tseng A, Sanchez KO, Shah H, Stanley BG, Gabriella I, Tseng A, Sanchez KO, Shah H & Stanley BG (2022). Stimulation of GABA Receptors in the Lateral Septum Rapidly Elicits Food Intake and Mediates Natural Feeding. Brain Sci 12, 848.

Gallagher JP & Hasuo H (1989). BicucullineL and phaclofenLsensitive components of NLmethylLDLaspartateLinduced hyperpolarizations in rat dorsolateral septal nucleus neurones. J Physiol 418, 367–377.

Gao X (2009). Electrophysiological effects of MCH on neurons in the hypothalamus. Peptides (NY) 30, 2025–2030.

Gavioli EC, Canteras NS & de Lima TCM (1999). Anxiogenic-like effect induced by substance P injected into the lateral septal nucleus. Neuroreport 10, 3399–3403.

Gavioli EC, Canteras NS & De Lima TCM (2002). The role of lateral septal NK1 receptors in mediating anxiogenic effects induced by intracerebroventricular injection of substance P. Behav Brain Res 134, 411–415.

Georgescu D, Sears RM, Hommel JD, Barrot M, Bolan CA, Marsh DJ, Bednarek MA, Bibb JA, Maratos-flier E, Nestler EJ & Dileone RJ (2005). The Hypothalamic Neuropeptide Melanin-Concentrating Hormone Acts in the Nucleus Accumbens to Modulate Feeding Behavior and Forced-Swim Performance. J Neurosci 25, 2933–2940.

Harthoorn L, Sañé A, Nethe M & van Heerikhuize J (2005). Multi-transcriptional profiling of melanin-concentrating hormone and orexin-containing neurons. Cell Mol Neurobiol 25, 1209–1223.

Hawes B, Kil E, Green B, O’Neill K, Fried S & Graziano M (2000). The melanin-concentrating hormone receptor couples to multiple G proteins to activate diverse intracellular signaling pathways. Endocrinology 141, 4524–4532.

Heldt SA & Ressler KJ (2007). Forebrain and midbrain distribution of major benzodiazepine-sensitive GABAA receptor subunits in the adult C57 mouse as assessed with in situ hybridization. Neuroscience 150, 370–385.

Hervieu G, Cluderay J, Harrison D, Meakin J, Maycox P, Nasir S & Leslie R (2000). The distribution of the mRNA and protein products of the melanin-concentrating hormone (MCH) receptor gene, slc-1, in the central nervous system of the rat. Eur J Neurosci 12, 1194–1216.

Hill J, Duckworth M, Murdock P, Rennie G, Sabido-David C, Ames RS, Szekeres P, Wilson S, Bergsma DJ, Gloger IS, Levy DS, Chambers JK & Muir AI (2001). Molecular Cloning and Functional Characterization of MCH2, a Novel Human MCH Receptor. J Biol Chem 276, 20125–20129.

Hörtnagl H, Tasan RO, Wieselthaler A, Kirchmair E, Sieghart W & Sperk G (2013). Patterns of mRNA and protein expression for 12 GABAA receptor subunits in the mouse brain. Neuroscience 236, 345–372.

Jego S, Glasgow SD, Herrera CG, Ekstrand M, Reed SJ, Boyce R, Friedman J, Burdakov D & Adamantidis AR (2013). Optogenetic identification of a rapid eye movement sleep modulatory circuit in the hypothalamus. Nat Neurosci 16, 1637–1643.

Kim T-K & Han P-L (2016). Physical Exercise Counteracts Stress-induced Upregulation of Melanin-concentrating Hormone in the Brain and Stress-induced Persisting Anxiety-like Behaviors. Exp Neurobiol 25, 163–173.

Kokkotou E, Jeon JY, Wang X, Marino FE, Carlson M, Trombly DJ, Maratos-flier E, Jeon JY, Wang X, Marino FE, Carlson M, Trombly DJ & Mice EM (2005). Mice with MCH ablation resist diet-induced obesity through strain-specific mechanisms. Am J Physiol Regul Integr Comp Physiol 02215, 117–124.

Kovacs EG, Szalay F & Halasy K (2005). Fasting-induced changes of neuropeptide immunoreactivity in the lateral septum of male rats. Acta Biol Hung 56, 185–197.

Krimer LS, Jakab RL & Goldman-Rakic PS (1997). Quantitative three-dimensional analysis of the catecholaminergic innervation of identified neurons in the macaque prefrontal cortex. J Neurosci 17, 7450–7461.

Lambe EK, Krimer LS & Goldman-Rakic PS (2000). Differential postnatal development of catecholamine and serotonin inputs to identified neurons in prefrontal cortex of rhesus monkey. J Neurosci 20, 8780–8787.

Lee V & Maguire J (2014). The impact of tonic GABAA receptor-mediated inhibition on neuronal excitability varies across brain region and cell type. Front Neural Circuits 8, 1–27.

Lembo PMC, Grazzini E, Cao J, Hubatsch DA, Pelletier M, Hoffert C, St-Onge S, Pou C, Labrecque J, Groblewski T, O’Donnell D, Payza K, Ahmad S & Walker P (1999). The receptor for the orexigenic peptide melanin-concentrating hormone is a G-protein-coupled receptor. Nat Cell Biol 1, 267–271.

Lin YF, Angelotti TP, Dudek EM, Browning MD & Macdonald RL (1996a). Enhancement of recombinant α1β1γ2L γ-aminobutyric acid(A) receptor whole-cell currents by protein kinase C is mediated through phosphorylation of both β1 and γ2L subunits. Mol Pharmacol 50, 185–195.

Lin YF, Angelotti TP, Dudek EM, Browning MD & Macdonald RL (1996b). Enhancement of recombinant α1β1γ2L γ-aminobutyric acid(A) receptor whole-cell currents by protein kinase C is mediated through phosphorylation of both β1 and γ2L subunits. Mol Pharmacol 50, 185–195.

Lin YF, Browning MD, Dudek EM & Macdonald RL (1994). Protein kinase C enhances recombinant bovine α1β1γ2L GABAA receptor whole-cell currents expressed in L929 fibroblasts. Neuron 13, 1421–1431.

Lipo E, Asrat S, Huo W, Sol A, Fraser CS & Isberg RR (2022). 5’ Untranslated mRNA Regions Allow Bypass of Host Cell Translation Inhibition by Legionella pneumophila. Infect Immun 90, e0017922.

Liu J-J, Tsien RW & Pang ZP (2022). Hypothalamic melanin-concentrating hormone regulates hippocampus-dorsolateral septum activity. Nat Neurosci 25, 61–71.

Ludwig D, Tritos N, Mastaitis J, Kulkarni R, Kokkotou E, Elmquist J, Lowell B, Flier J & Maratos-Flier E (2001). Melanin-concentrating hormone overexpression in transgenic mice leads to obesity and insulin resistance. J Clin Invest 107, 379–386.

Mansour A, Fox CA, Burke S, Meng F, Thompson RC, Akil H & Watson SJ (1994). Mu, Delta, and Kappa Opioid Receptor mRNA Expression in the Rat CNS: An In Situ Hybridization Study. J Comp Neurol 350, 412–438.

le Merrer J, Cagniard B & Cazala P (2006). Modulation of anxiety by mu-opioid receptors of the lateral septal region in mice. Pharmacol Biochem Behav 83, 465–479.

Mickelsen L, Bolisetty M, Chimileski B, Fujita A, Beltrami E, Costanzo J, Naparstek J, Robson P & Jackson A (2019). Single-cell transcriptomic analysis of the lateral hypothalamic area reveals molecularly distinct populations of inhibitory and excitatory neurons. Nat Neurosci 22, 642–656.

Mickelsen LE, Kolling FW, IV, Chimileski BR, Fujita A, Norris C, Chen K, Nelson CE & Jackson AC (2017). Neurochemical Heterogeneity Among Lateral Hypothalamic Hypocretin/Orexin and Melanin-Concentrating Hormone Neurons Identified Through Single-Cell Gene Expression Analysis. eNeuro 4, 13–17.

Mogi K, Funabashi T, Mitsushima D, Hagiwara H & Kimura F (2005). Sex Difference in the Response of Melanin-Concentrating Hormone Neurons in the Lateral Hypothalamic Area to Glucose, as Revealed by the Expression of Phosphorylated Cyclic Adenosine 3,5-Monophosphate Response Element-Binding Protein. Endocrinology 146, 3325–3333.

Monzon M, de Souza M, Izquierdo L, Izquierdo I, Barros D & de Barioglio S (1999). Melanin-concentrating hormone (MCH) modifies memory retention in rats. Peptides (NY) 20, 1517– 1519.

Mu M-D, Geng H-Y, Rong K-L, Peng R-C, Wang S-T, Geng L-T, Qian Z-M, Yung W-H & Ke Y (2020). A limbic circuitry involved in emotional stress-induced grooming. Nat Commun 11, 2261.

Mul JD, la Fleur SE, Toonen PW, Afrasiab-Middelman A, Binnekade R, Schetters D, Verheij MMM, Sears RM, Homberg JR, Schoffelmeer ANM, Adan RAH, DiLeone RJ, de Vries TJ & Cuppen E (2011). Chronic loss of melanin-concentrating hormone affects motivational aspects of feeding in the rat. PLoS One 6, e19600.

Mystkowski P, Seeley RJ, Hahn TM, Baskin DG, Havel PJ, Matsumoto AM, Wilkinson CW, Peacock-Kinzig K, Blake KA & Schwartz MW (2000). Hypothalamic Melanin-Concentrating Hormone and Estrogen-Induced Weight Loss. J Neurosci 20, 8637–8642.

Negishi K, Payant MA, Schumacker KS, Wittmann G, Butler RM, Lechan RM, Steinbusch HWM, Khan AM & Chee MJ (2020). Distributions of hypothalamic neuron populations coexpressing tyrosine hydroxylase and the vesicular GABA transporter in the mouse. J Comp Neurol 528, 1833–1855.

Noble E, Hahn J, Konanur V, Hsu T, Page S, Cortella A, Liu C, Song M, Suarez A, Szujewski C, Rider D, Clarke J, Darvas M, Appleyard S & Kanoski S (2018). Control of Feeding Behavior by Cerebral Ventricular Volume Transmission of Melanin-Concentrating Hormone. Cell Metab 28, 55–68.e7.

Noguchi T, Makino S, Shinahara M, Nishiyama M, Hashimoto K & Terada Y (2013). Effects of gold thioglucose treatment on central corticotrophin-releasing hormone systems in mice. J Neuroendocrinol 25, 340–349.

Parfitt GM, Nguyen R, Yoon Bang J, Aqrabawi AJ, Tran MM, Seo K, Richards BA & Kim JC (2017). Bidirectional Control of Anxiety-Related Behaviors in Mice: Role of Inputs Arising from the Ventral Hippocampus to the Lateral Septum and Medial Prefrontal Cortex. Neuropsychopharmacology 42, 1715–1728.

Pissios P, Bradley R & Maratos-Flier E (2006). Expanding the scales: The multiple roles of MCH in regulating energy balance and other biological functions. Endocr Rev 27, 606–620.

Pissios P, Trombly DJ, Tzameli I & Maratos-Flier E (2003). Melanin-concentrating hormone receptor 1 activates extracellular signal-regulated kinase and synergizes with Gs-coupled pathways. Endocrinology 144, 3514–3523.

Poisbeau P, Cheney MC, Browning MD & Mody I (1999). Modulation of synaptic GABA(A) receptor function by PKA and PKC in adult hippocampal neurons. J Neurosci 19, 674–683.

Qu D, Ludwig D, Gammeltoft S, Piper M, Pelleymounter M, Cullen M, Mathes W, Przypek R, Kanarek R & Maratos-Flier E (1996). A role for melanin-concentrating hormone in the central regulation of feeding behaviour. Nature 380, 243–247.

Rao Y, Lu M, Ge F, Marsh DJ, Qian S, Wang AH, Picciotto MR & Gao XB (2008). Regulation of synaptic efficacy in hypocretin/orexin-containing neurons by melanin concentrating hormone in the lateral hypothalamus. J Neurosci 28, 9101–9110.

Risold PY & Swanson LW (1997a). Connections of the rat lateral septal complex. Brain Res Brain Res Rev 24, 115–195.

Risold PY & Swanson LW (1997b). Chemoarchitecture of the rat lateral septal nucleus. Brain Res Brain Res Rev 24, 91–113.

Rondini TA, de Crudis Rodrigues B, de Oliveira AP, Bittencourt JC & Elias CF (2007). Melanin-concentrating hormone is expressed in the laterodorsal tegmental nucleus only in female rats. Brain Res Bull 74, 21–28.

Rossi M, Choi SJ, O’Shea D, Miyoshi T, Ghatei MA & Bloom SR (1997). Melanin-concentrating hormone acutely stimulates feeding, but chronic administration has no effect on body weight. Endocrinology 138, 351–355.

Ruiz-Viroga V, Urbanavicius J, Torterolo P & Lagos P (2021). In vivo uptake of a fluorescent conjugate of melanin-concentrating hormone in the rat brain. J Chem Neuroanat 114, 101959.

Saito Y, Cheng M, Leslie FM & Civelli O (2001). Expression of the melanin-concentrating hormone (MCH) receptor mRNA in the rat brain. J Comp Neurol 435, 26–40.

Saito Y, Nothacker HP, Wang Z, Lin SHS, Leslie F & Civelli O (1999). Molecular characterization of the melanin-concentrating-hormone receptor. Nature 400, 265–269.

Sears RM, Liu RJ, Narayanan NS, Sharf R, Yeckel MF, Laubach M, Aghajanian GK & DiLeone RJ (2010). Regulation of nucleus accumbens activity by the hypothalamic neuropeptide melanin-concentrating hormone. J Neurosci 30, 8263–8273.

Sheehan TP, Chambers RA & Russell DS (2004). Regulation of affect by the lateral septumL: implications for neuropsychiatry. Brain Res Brain Res Rev 46, 71–117.

Shimada M, Tritos NA, Lowell BB, Flier JS & Maratos-Flier E (1998). Mice lacking melanin-concentrating hormone are hypophagic and lean. Nature 396, 670–674.

Skofitsch G, Jacobowitz DM & Zamir N (1985). Immunohistochemical Localization of a Melanin Concentrating Hormone-Like Peptide in the Rat Brain. Brain Res Bull 15, 635– 649.

Smith D, Davis R, Rorick-Kehn L, Morin M, Witkin J, McKinzie D, Nomikos G & Gehlert D (2006). Melanin-concentrating hormone-1 receptor modulates neuroendocrine, behavioral, and corticolimbic neurochemical stress responses in mice. Neuropsychopharmacology 31, 1135–1145.

Stengel A & Taché Y (2014). CRF and urocortin peptides as modulators of energy balance and feeding behavior during stress. Front Neurosci 8, 1–10.

Swanson LW & Cowan WM (1977). An autoradiographic study of the organization of the efferet connections of the hippocampal formation in the rat. J Comp Neurol 172, 49–84.

Takase K, Kikuchi K, Tsuneoka Y, Oda S & Kuroda M (2014). Meta-Analysis of Melanin-Concentrating Hormone Signaling-Deficient Mice on Behavioral and Metabolic Phenotypes. PLoS One 9, 99961.

Tan CP et al. (2002). Melanin-concentrating hormone receptor subtypes 1 and 2: Species-specific gene expression. Genomics 79, 785–792.

Teixeira PDS, Wasinski F, Lima LB, Frazão R, Bittencourt JC & Donato J (2020). Regulation and neurochemical identity of melanin-concentrating hormone neurones in the preoptic area of lactating mice. J Neuroendocrinol 32, e12818.

Terrill SJ, Subramanian KS, Lan R, Liu CM, Cortella AM, Noble EE & Kanoski SE (2020). Nucleus accumbens melanin-concentrating hormone signaling promotes feeding in a sex-specific manner HHS Public Access. Neuropharmacology 178, 108270.

Valeyev AY, Hackman JC, Holohean AM, Wood PM, Katz JL & Davidoff RA (1999). GABA- induced Cl- current in cultured embryonic human dorsal root ganglion neurons. J Neurophysiol 82, 1–9.

Van Pett K, Viau V, Bittencourt J, Chan R, Li H, Arias C, Prins G, Perrin M, Vale W & Sawchenko P (2000). Distribution of mRNAs encoding CRF receptors in brain and pituitary of rat and mouse. J Comp Neurol 428, 191–212.

Verret L, Goutagny R, Fort P, Cagnon L, Salvert D, Léger L, Boissard R, Salin P, Peyron C & Luppi P-H (2003). A role of melanin-concentrating hormone producing neurons in the central regulation of paradoxical sleep. BMC Neurosci 4, 19.

Wang C & Kotz C (2002). Urocortin in the lateral septal area modulates feeding induced by orexin A in the lateral hypothalamus. Am J Physiol Regul Integr Comp Physiol 283, 358– 367.

Wu M, Dumalska I, Morozova E, van den Pol A & Alreja M (2009). Melanin-concentrating hormone directly inhibits GnRH neurons and blocks kisspeptin activation, linking energy balance to reproduction. Proc Natl Acad Sci U S A 106, 17217–17222.

Xu Y, Lu Y, Cassidy R, Mangieri L, Zhu C, Huang X, Jiang Z, Justice N, Xu Y, Arenkiel B & Tong Q (2019). Identification of a neurocircuit underlying regulation of feeding by stress-related emotional responses. Nat Commun 10, 3446.

Zhao C, Eisinger B & Gammie SC (2013). Characterization of GABAergic Neurons in the Mouse Lateral Septum: A Double Fluorescence In Situ Hybridization and Immunohistochemical Study Using Tyramide Signal Amplification. PLoS One 8, e73750.

Zheng H, Patterson LM, Morrison C, Banfield BW, Randall JA, Browning KN, Travagli RA & Berthoud HR (2005). Melanin concentrating hormone innervation of caudal brainstem areas involved in gastrointestinal functions and energy balance. Neuroscience 135, 611–625.

