## Supplementary figures and images for "Inhibitory actions of melanin-concentrating hormone in the lateral septum"

### Supporting Figure 1

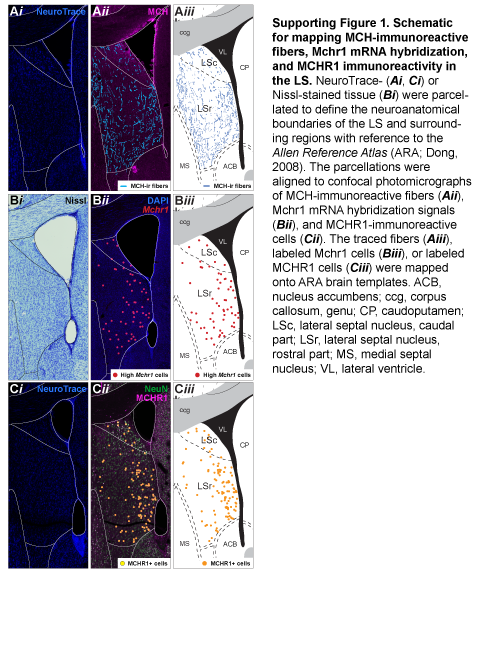

### Supporting Figure 2

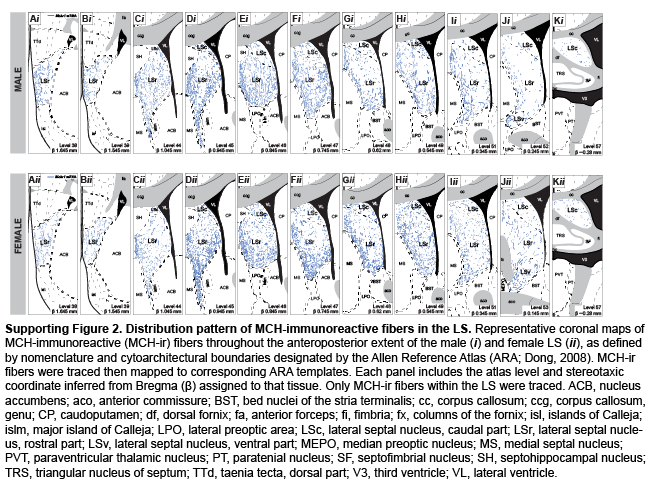

### Supporting Figure 3

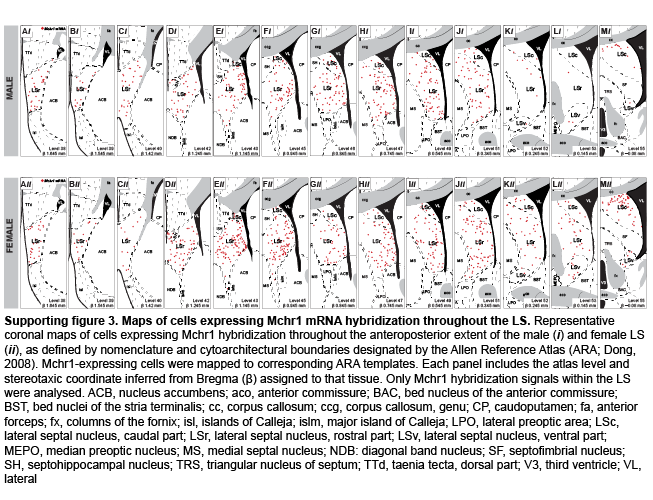

### Supporting Figure 4

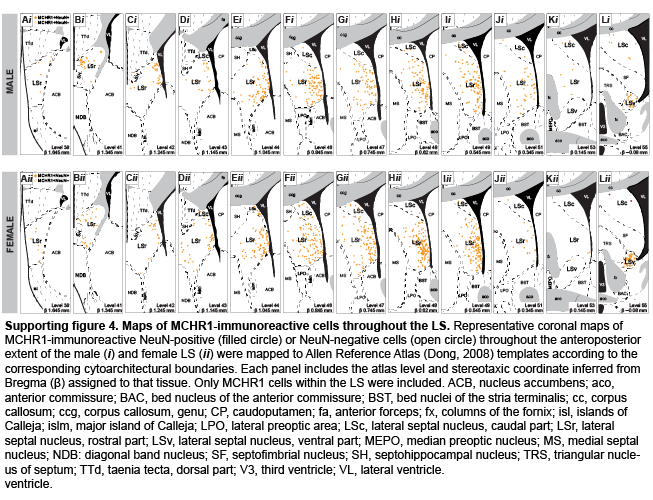
